# Impact of cannabis use on immune cell populations and the viral reservoir in people with HIV on suppressive antiretroviral therapy

**DOI:** 10.1101/2022.12.22.521628

**Authors:** Shane D. Falcinelli, Alicia D Cooper-Volkheimer, Lesia Semenova, Ethan Wu, Alexander Richardson, Manickam Ashokkumar, David M Margolis, Nancie M. Archin, Cynthia D Rudin, David Murdoch, Edward P Browne

**Affiliations:** Department of Medicine, University of North Carolina at Chapel Hill, Chapel Hill, North Carolina, United States of America; Department of Microbiology and Immunology, University of North Carolina at Chapel Hill, Chapel Hill, North Carolina, United States of America; HIV Cure Center, University of North Carolina at Chapel Hill, Chapel Hill, North Carolina, United States of America; Department of Medicine, Duke University, Durham, North Carolina, United States of America; Department of Computer Science, Duke University, Durham, North Carolina, United States of America

## Abstract

HIV infection remains incurable due to the persistence of a viral reservoir during antiretroviral therapy. Cannabis (CB) use is prevalent amongst people with HIV (PWH), but the impact of CB on the latent HIV reservoir has not been investigated. Peripheral CD4 and CD8 T cells from a cohort of CB-using PWH and a matched cohort of non-users on antiretroviral therapy were evaluated for expression of maturation/activation markers, HIV-specific T cell responses, and the frequency of intact proviral DNA. CB use was associated with increased abundance of naïve T cells, reduced effector T cells, and reduced expression of activation markers. CB users also exhibited reduced levels of exhausted and senescent T cells compared to non-using controls. HIV-specific CD8 T cell responses were unaffected by CB use. While the abundance of intact proviruses was not significantly affected by CB use across the whole cohort, we observed that, for participants with high frequency of NKG2A or CD16 expression in NK cells, CB use was associated with a smaller intact HIV reservoir. This analysis is consistent with the hypothesis that CB use reduces activation, exhaustion and senescence in the T cells of PWH and may influence the size of the HIV reservoir.

## Introduction

40 million people worldwide are living with human immunodeficiency virus (HIV) infection (1). Although infection can be effectively treated with antiretroviral therapy (ART), the persistence of latently infected reservoir cells requires lifelong ART and prevents an HIV cure (2–4). The overall size of this reservoir declines slowly over time, but at a rate that precludes natural elimination of the reservoir within a human lifetime (5, 6). The reservoir is also highly dynamic, with temporal waxing and waning of individual proviruses within T cell clones during ART among other phenomena (7, 8).

Untreated HIV infection is associated with greatly elevated immune activation and progressive depletion of CD4 T cells, along with exhaustion of cytotoxic lymphocytes (CTLs) (9, 10). This immune activation is a strong predictor of disease progression (11), but the mechanisms that contribute to it are not fully understood (12). ART helps to mitigate this immune activation and prevents the development of Acquired Immune Deficiency Syndrome (AIDS), but even on therapy, people with HIV (PWH) experience elevated levels of immune activation and inflammation (13). In particular, ART-suppressed PWH exhibit a number of alterations to the abundance and phenotype of immune subsets, including an increased proportion of effector cells, elevated expression of activation markers, and elevated concentrations of pro-inflammatory cytokines (14, 15). These perturbations lead to an overall state of elevated immune exhaustion and senescence in ART-suppressed PWH. This chronic immune activation is likely to contribute to increased morbidity amongst ART-treated PWH compared to uninfected people, including lung, kidney, liver and cardiovascular disease (16). The driving mechanisms of this elevated inflammation and immune disfunction during ART are unclear but may involve stimulation of the innate immune system by residual proviral DNA/RNA, antigen-specific responses to cells that contain translation-competent proviruses, or long-lasting immune system alterations induced during acute or untreated chronic infection, such as damage to intestinal permeability and microbial translocation (17).

Cannabis (CB) use is prevalent among PWH, with up to 49% percent of PWH reporting some use (18–20). Despite its high medicinal and recreational use and increasing legality, little is known about its effect on the immune system. Δ - 9-tetrahydrocannabinol (THC), the major psychoactive cannabinoid constituent, exerts its effects via two G-protein coupled receptors CB_1_ and CB_2_. CB_1_ is largely present in brain and neurons and mediates the psychoactive properties of cannabis. CB_2_, by contrast, is abundantly expressed in immune cells, including CD4 T cells (21–24). At the molecular level, activation of cannabinoid receptors leads to complex pleotropic cell-type and context-dependent cellular signaling effects. Prominent among these effects are modulation of the MAPK, JNK, and PI3K/AKT pathways (25–27). Transcriptomic analyses of human immune cells have revealed that cannabis induces numerous transcriptomic pathway alterations in CD4 T cells (28). Currently, cannabinoids are canonically thought of as immunosuppressive, reducing TNF-α production, T cell proliferation and IL-2 expression after TCR-mediated activation (25, 29). Other data suggest THC can inhibit T-cell dependent antigen immune responses and skew CD4 T cells towards a Th_2_ phenotype (30).

Even less is known regarding the virological effects of cannabis use within the context of HIV infection. *In vitro* studies have shown inhibition of HIV replication in cultured cells by synthetic cannabinoids (31), and some clinical studies suggest reduced viral loads in PWH using cannabis (32). The impact of cannabis use on the immune system of PWH has been somewhat controversial. In some studies, cannabis has been reported to lower inflammation markers and immune activation in ART-suppressed PWH (33–37), while other studies have found either no major impact of CB use on the immune system of PWH or an increase in inflammatory markers (38–40). Animal studies of CB administration and HIV infection have also yielded contradictory findings: THC dosing of HIV-infected huPBL-SCID mice increased viral loads, while dosing of SIV-infected Rhesus Macaques exhibited decreased mortality and viral loads (41, 42).

Therefore, further investigation is needed to fully assess the impact of cannabis use on viral dynamics and on the HIV reservoir. However, quantification of the fraction of the reservoir capable of re-establishing infection upon ART cessation is complicated by a predominance of defective proviruses with inactivating deletions or hypermutation that renders the virus unable to replicate (43, 44). To address this, the intact proviral DNA assay (IPDA) was developed to more accurately quantify HIV DNA that is likely to be replication-competent (45).

In this study, we performed simultaneous high dimensional immunophenotyping and HIV reservoir quantification using IPDA on a CB-using cohort of PWH on suppressive ART to examine the impact of chronic cannabis use on cellular activation, immune response, and viral reservoir size. We observed clear alterations of key immune cell subsets and changes in T cell activation and senescence in CB users. Furthermore, we observed a trend towards a smaller HIV reservoir size in subgroups of CB users, suggesting that the formation and/or maintenance of the HIV reservoir may be impacted by CB use.

## Results

### Cannabis using PWH cohort

To study the impact of cannabis (CB) use on immune cell phenotype/function and on the HIV reservoir, we recruited a cohort of 33 CB-using people with HIV (PWH) and 42 non-using PWH as controls. All participants had been HIV infected for more than 2 years (median 12 years, range 2-34 years) and were ART-suppressed (<50 RNA copies/mL) for at least 12 months (median 10 years, range 2-28 years). Cannabis users were defined by greater than 12 days of self-reported CB use within the last 90 days and a positive urine test, while non-users were defined as having no reported CB or other illicit drug use in the last 12 months and a negative urine test. Individuals with recent or previous regular use of drugs other than cannabis, as well as individuals with alcohol use disorder, were excluded. Relevant clinical and demographic information for the cohorts is shown in **Table 1**. Overall, the two groups had similar characteristics in terms of age, sex, race, years of infection and therapy, and CD4 nadir.

**Table 1:**
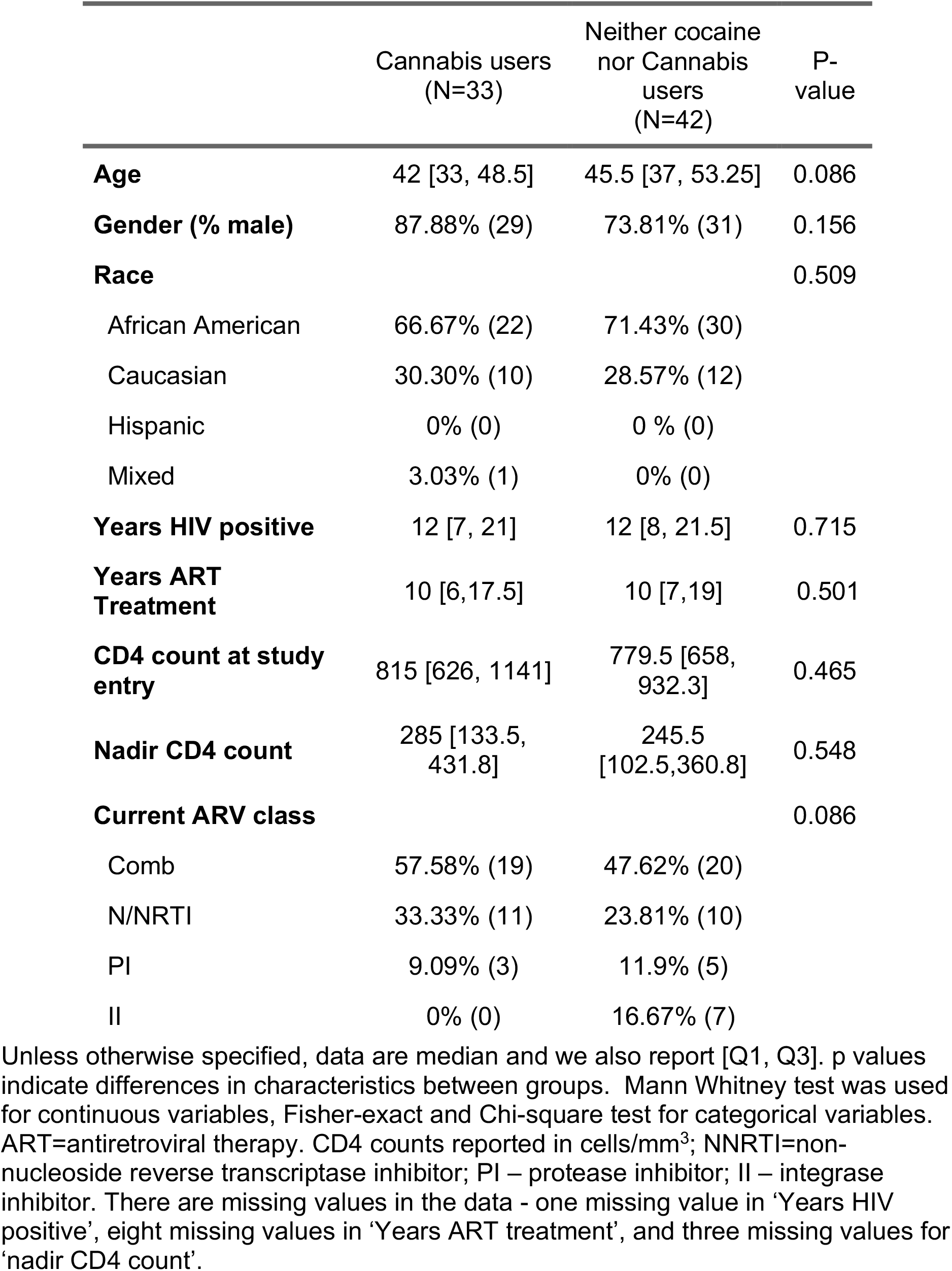
Participant demographic and clinical characteristics

|  | Cannabis users<br>(N=33) | Neither cocaine<br>nor Cannabis<br>users<br>(N=42) | P-value |
| --- | --- | --- | --- |
| **Age** | 42 [33, 48.5] | 45.5 [37, 53.25] | 0.086 |
| **Gender (% male)** | 87.88% (29) | 73.81% (31) | 0.156 |
| **Race** |  |  | 0.509 |
| African American | 66.67% (22) | 71.43% (30) |  |
| Caucasian | 30.30% (10) | 28.57% (12) |  |
| Hispanic | 0% (0) | 0 % (0) |  |
| Mixed | 3.03% (1) | 0% (0) |  |
| **Years HIV positive** | 12 [7, 21] | 12 [8, 21.5] | 0.715 |
| **Years ART**<br>**Treatment** | 10 [6,17.5] | 10 [7,19] | 0.501 |
| **CD4 count at study**<br>**entry** | 815 [626, 1141] | 779.5 [658,<br>932.3] | 0.465 |
| **Nadir CD4 count** | 285 [133.5,<br>431.8] | 245.5<br>[102.5,360.8] | 0.548 |
| **Current ARV class** |  |  | 0.086 |
| Comb | 57.58% (19) | 47.62% (20) |  |
| N/NRTI | 33.33% (11) | 23.81% (10) |  |
| PI | 9.09% (3) | 11.9% (5) |  |
| II | 0% (0) | 16.67% (7) |  |

Unless otherwise specified, data are median and we also report [Q1, Q3]. p values indicate differences in characteristics between groups. Mann Whitney test was used for continuous variables, Fisher-exact and Chi-square test for categorical variables. ART=antiretroviral therapy. CD4 counts reported in cells/mm<sup>3</sup>; NNRTI=non-nucleoside reverse transcriptase inhibitor; PI – protease inhibitor; II – integrase inhibitor. There are missing values in the data - one missing value in 'Years HIV positive', eight missing values in 'Years ART treatment', and three missing values for 'nadir CD4 count'.

### CB use causes alterations to T cell subset abundance in PWH

To determine the impact of CB use on the immune cells of a PWH population, we performed high dimensional flow cytometry immunophenotyping designed to differentiate T cell subsets, natural killer (NK), and NK T cells (NKT) (**Figure 1A**). T cells, NK cells and NKT were identified based on CD3 and CD56 expression: T cells defined by CD3^+^/CD56^-^, NK cells defined by CD3^-^/CD56^+^, and NKT cells defined by CD3^+^/CD56^+^. T cells were then further subclassified as CD4 T or CD8 T based on CD4 or CD8 expression respectively. Markers of cellular senescence (KLRG1), exhaustion (PD-1), and cellular activation (HLA-DR, CD38) were assessed on T cell, NK, and NKT sub-populations. Comparing the CB-using cohort to the non-user controls, we observed no significant differences for overall abundance of CD4 T cells, CD8T cells, NK cells, or NKT cells between the cohorts (**Figure 1B, 1C**). Within the CD4 and CD8 T cell compartments, we then examined the abundance of T cells within broad developmental subtypes. We used CCR7 and CD45RA expression to assign cells as naïve (Tn - CD45RA^+^/CCR7^+^), central memory (Tcm - CD45RA^-^/CCR7^+^), effector memory (Tem - CD45RA^-^/CCR7^-^) or terminally differentiated effector cells (Teff - CD45RA^+^/CCR7^-^) (**Figure 1A**). We then compared the relative frequencies of these subsets between the CB using and non-using cohorts (**Figure 2A, 2B**). For CD4 T cells, we observed a trend towards a larger naïve compartment in CB users compared to non-users (median 36.3% vs. 27.0%, p = 0.056 Mann Whitney U test) and no significant change to the fraction of Tcm, Tem and Teff cells. For CD8 T cells, we observed a significantly increased fraction of naïve cells for CB users (median 38.6% vs 26.2%, p = 0.004, Mann-Whitney U test), no change to Tcm and Tem populations, and a significantly smaller fraction of Teff cells for CB users (median 27.6% vs 33.2, p = 0.013, Mann-Whitney U test). Overall, these data demonstrate that CB use causes significant alterations to the fraction of CD4 and CD8 T cells within specific memory and effector subsets for PWH on suppressive ART.

**Figure 1.**
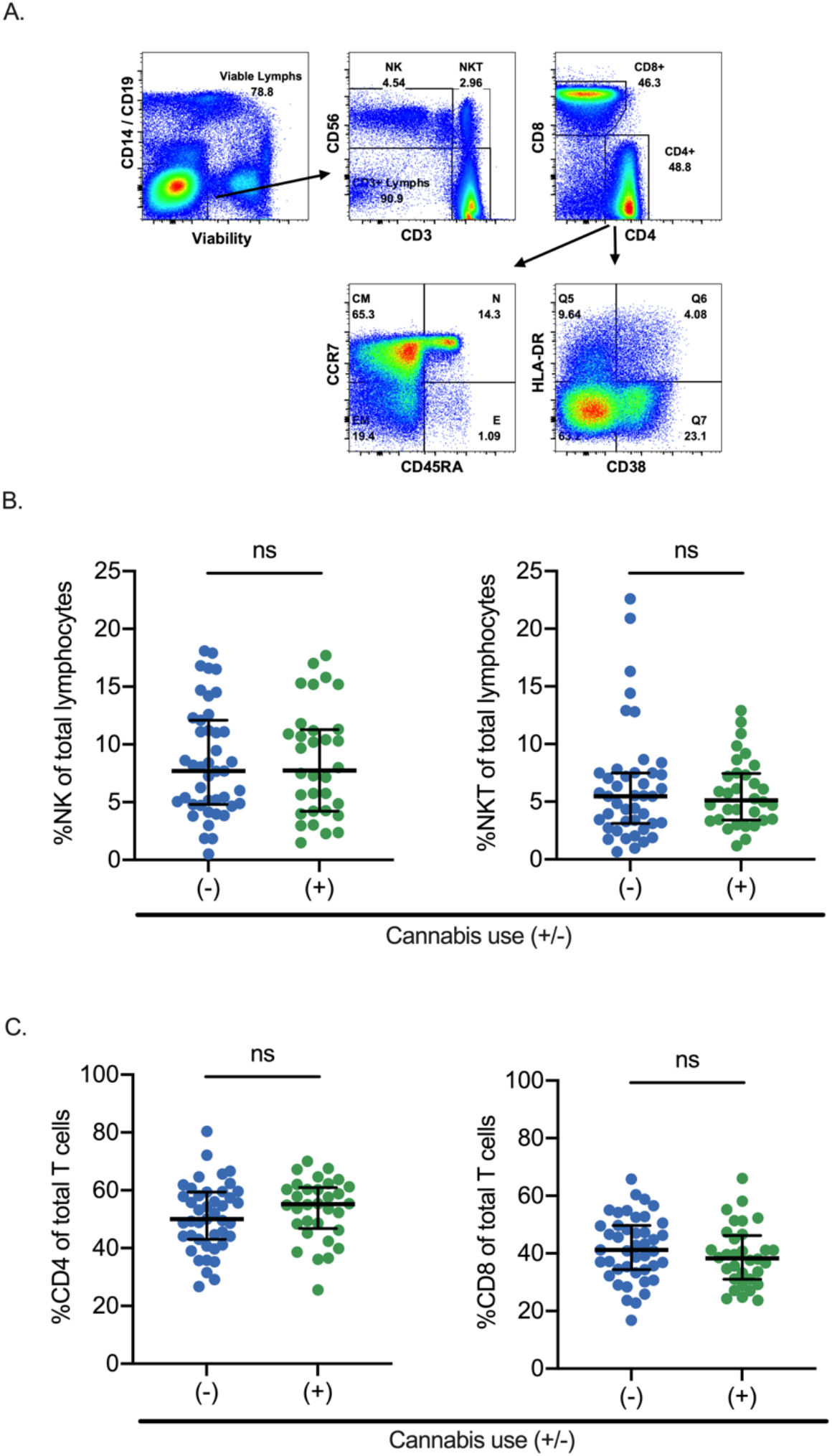
Impact of cannabis use on T cell, NK and NKT cell populations in people with HIV. **A**. Representative flow cytometry gating for NK cells, NKT cells, CD4 T and CD8 T cells. N=naïve, CM=central memory, EM=effector memory, E=terminally differentiated effector cells. **B**. NK (CD56^+^, CD3^-^) and NKT cells (CD56^+^, CD3^+^) are expressed as a fraction of the total live lymphocyte population. **C**. CD4 and CD8 T cells are expressed as a fraction of the T cell (CD3^+^, CD56^-^) population. Each dot represents an individual participant. Statistical significance determined using a Mann-Whitney U test. ns = not significant (p < 0.05). Median and IQR are displayed.

**Figure 2:**
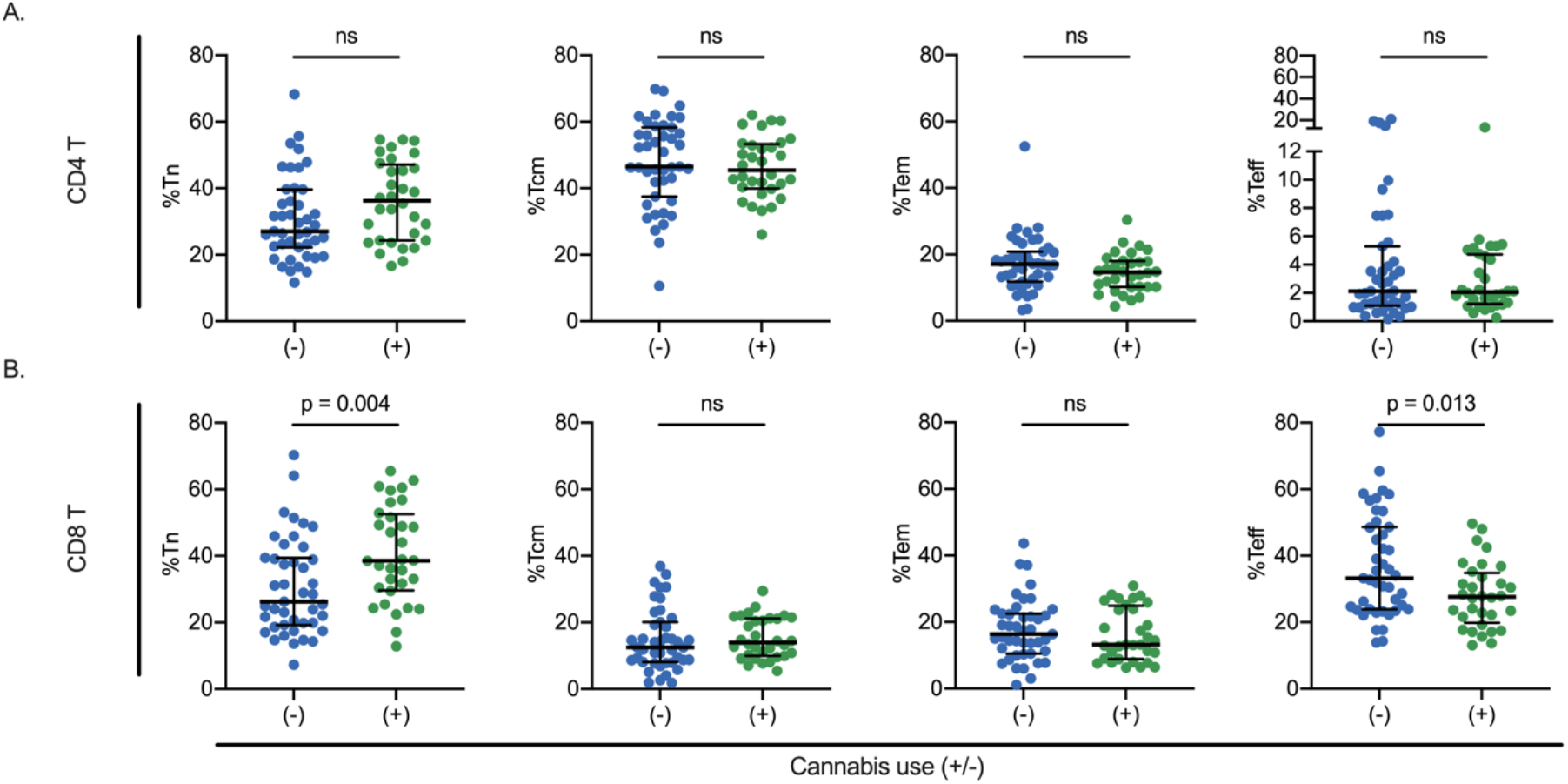
Impact of cannabis use on T cell subset proportions in people with HIV. Proportional abundance of T cells in naïve (Tn), central memory (Tcm), effector memory (Tem) and effector (Teff) subsets for CD4 T cells **(A)** and CD8 T cells **(B)**. Cell subsets assigned by expression of the surface markers CD45RA and CCR7 – Tn:CD45RA^+^/CCR7^+^, Tcm:CD45RA^-^/CCR7^+^, Tem:CD45RA^-^/CCR7^-^, Teff:CD45RA^+^/CCR7^-^. Each dot represents an individual participant. Statistical significance determined using a Mann-Whitney test. P values for comparisons where CB users were significantly different (p < 0.05) to non-user controls are displayed. Ns = not significant (p ≥ 0.05). Median and IQR are displayed for each dataset.

### CB use is associated with reduced expression of T cell activation markers

CD38 and HLA-DR, two of the most studied markers of immune activation, are of prognostic importance in HIV infection (46, 47). Previous reports have indicated that CB use can reduce T cell activation and inflammation in PWH (48–50). To examine this possibility, we examined the impact of CB use on the proportion of T cells that express activation and exhaustion markers. We first compared expression of three markers (HLA-DR, CD38 and PD-1) in total CD4 T cells, as well as within individual CD4 T cell memory subtypes (**Figure 3**). Cannabis users exhibited significantly reduced surface expression of PD-1 in total CD4 T cells (median 45.0% vs 55.4%, p = 0.007, Mann-Whitney U test), with significantly lower expression in central and effector memory subsets (p = 0.047 and 0.015, respectively). We observed little difference in CD4 T cell HLA-DR expression between CB users and non-users. However, there was differential expression of CD38 within CD4 T cell memory subsets. Specifically, CB use was associated with reduced CD38 expression CD4 Tn, Tcm and Teff cells. These data indicate that CB use can have differential effects on CD4 T cells depending on the subpopulation and highlight the importance of considering individual memory subtypes in analysis. Overall, we conclude that CB use modulates expression of activation markers in multiple types of CD4 T cells.

**Figure 3.**
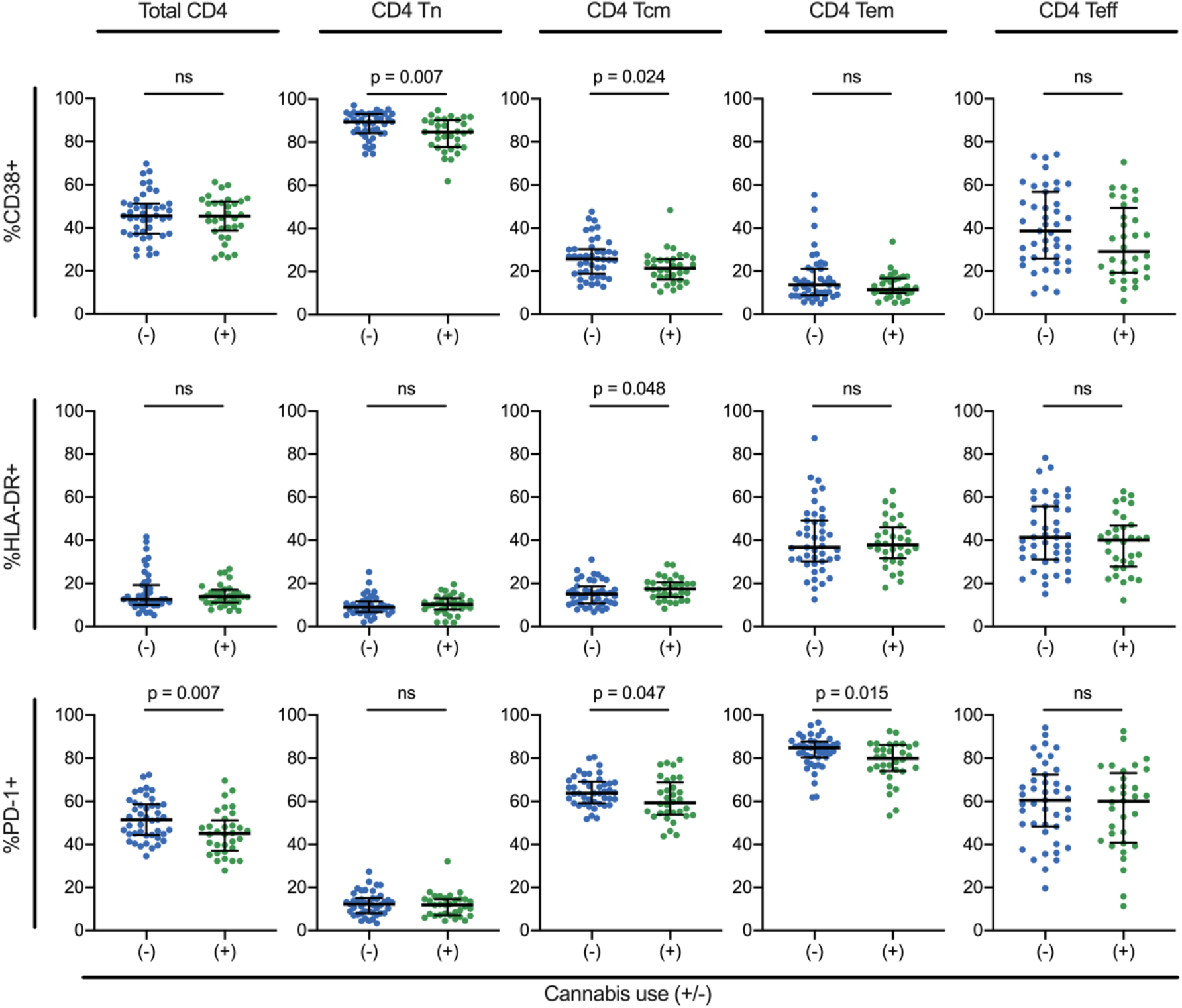
Impact of CB use on expression of activation markers for CD4 T cell subsets. Expression levels (percent positive) for three different immune activation markers (HLA-DR, CD38 and PD-1) are shown for total CD4 T cells and for specific subsets (Tn, Tcm, Tem, Teff). Each dot represents an individual participant. Statistical significance determined using a Mann-Whitney test. P values for comparisons where CB users were significantly different (p < 0.05) to non-user controls are displayed. Ns = not significant (p ≥ 0.05). Median and IQR are displayed for each dataset.

We also examined the expression of CD38, HLA-DR and PD-1 within total CD8 T cells across the cohorts (**Figure 4**). In addition to representing an activation marker, PD-1 is also expressed in a subset of exhausted CD8 T cells that accumulate during HIV infection (51). Although we observed no difference in PD-1 expression on total CD8 T cells between groups, we observed decreased expression of PD-1 in CB users within CD8 Tn and Tcm cells. Unlike CD4 T cells, CD38 expression was not significantly impacted by CB use in any CD8 T cell subset. Altogether, these data demonstrate a significant effect of CB use on the activation level for multiple CD4 and CD8 T cell subtypes that may influence the overall health of PWH and the immune response to HIV.

**Figure 4.**
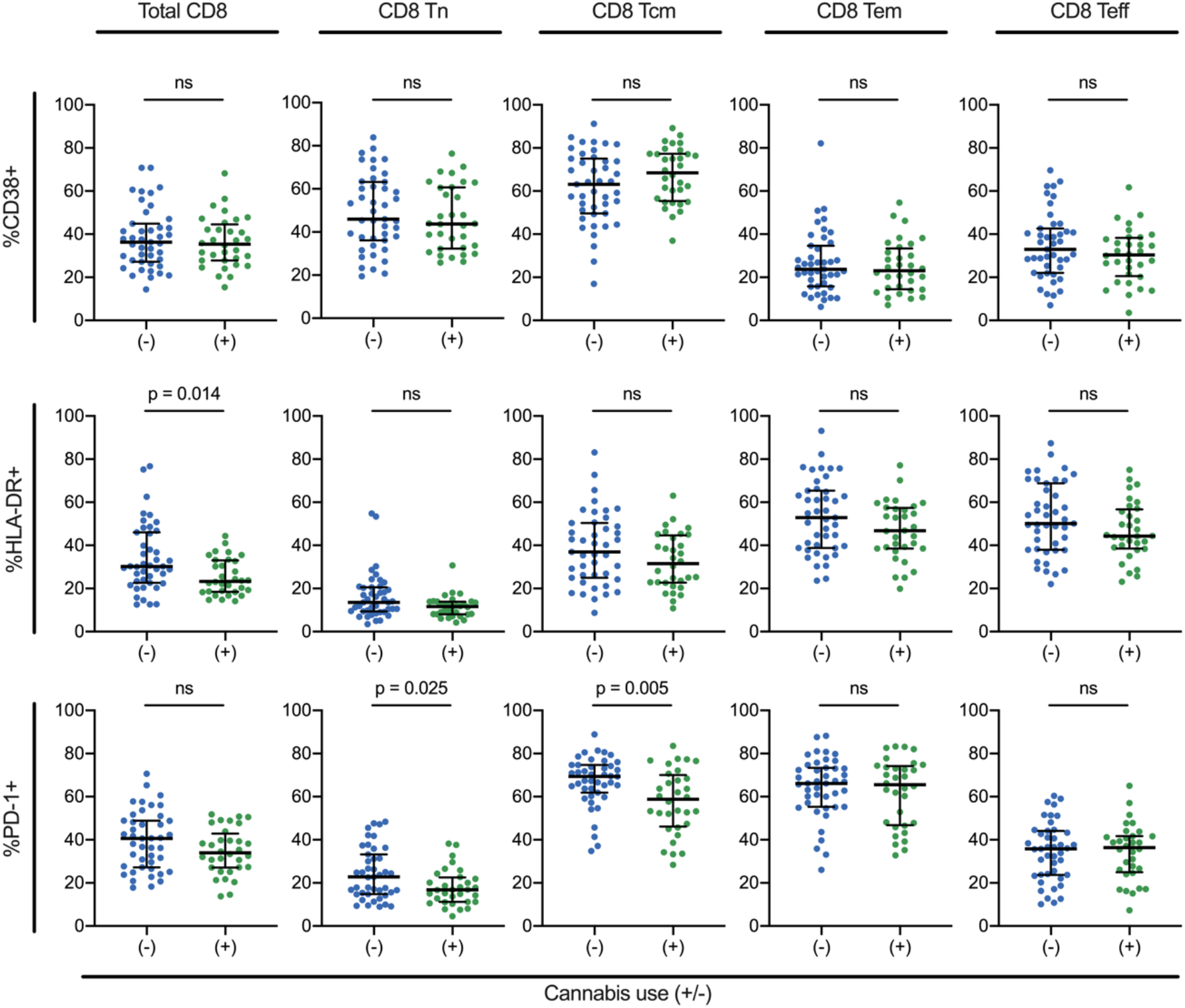
Impact of CB use on expression of activation markers for CD8 T cell subsets. Expression levels (percent positive) for three different immune activation markers (HLA-DR, CD38 and PD-1) are shown for total CD8 T cells and for specific subsets (Tn, Tcm, Tem, Teff). Each dot represents an individual participant. Statistical significance determined using a Mann-Whitney test. P values for comparisons where CB users were significantly different (p < 0.05) to non-user controls are displayed. Ns = not significant (p ≥ 0.05). Median and IQR are displayed for each dataset.

### CB use is associated with reduced immune senescence in the CD8 T cells of PWH

HIV infection promotes accelerated immune senescence, indicated by elevated expression of markers of immune senescence and exhaustion, even in individuals undergoing suppressive ART (9, 52, 53). A key marker of cellular exhaustion and senescence for HIV infection is killer cell lectin-like receptor subfamily G member 1 (KLRG1) (54, 55). Previous data have shown that, under conditions of chronic antigen stimulation, CD8 T cells become increasingly dysfunctional and upregulate expression of KLRG1 (56). Human T cells expressing KLRG1 exhibit significantly attenuated proliferation potential (57) and KLRG1 blockade can restore function to HIV-specific CD8 T cells (58). Therefore, in order to assess the impact of CB-use on immune senescence, KLRG1 expression levels between the CB-using and non-using cohorts across different T cell subtypes were compared. While there was no difference in KLRG1 expression on total CD4 T cells (**Figure 5A**), we observed decreased expression in total CD8 T cells for CB users compared to non-users (**Figure 5B**, median 48.9% versus 68.3% respectively, p < 0.001, Mann-Whitney U test). We then looked further into the impact of CB use on KLRG1 expression in individual CD4 and CD8 T cell subsets. As expected, naïve CD4 and CD8 T cells expressed the lowest levels of KLRG1, followed by Tcm. By contrast, Tem and Teff exhibited higher levels of KLRG1 expression. CB use had no significant impact on KLRG1 expression levels in any of the major CD4 T cell maturation subtypes (Tn, Tcm, Tem, Teff). By contrast, all CD8 T cells subsets except Tcm exhibited a significant reduction in KLRG1 levels in CB users compared to non-users. These data indicate that CB use is associated with reduced CD8 T cell exhaustion and senescence in the context of treated HIV infection, but that CB use has a more limited impact on CD4 T cells.

**Figure 5.**
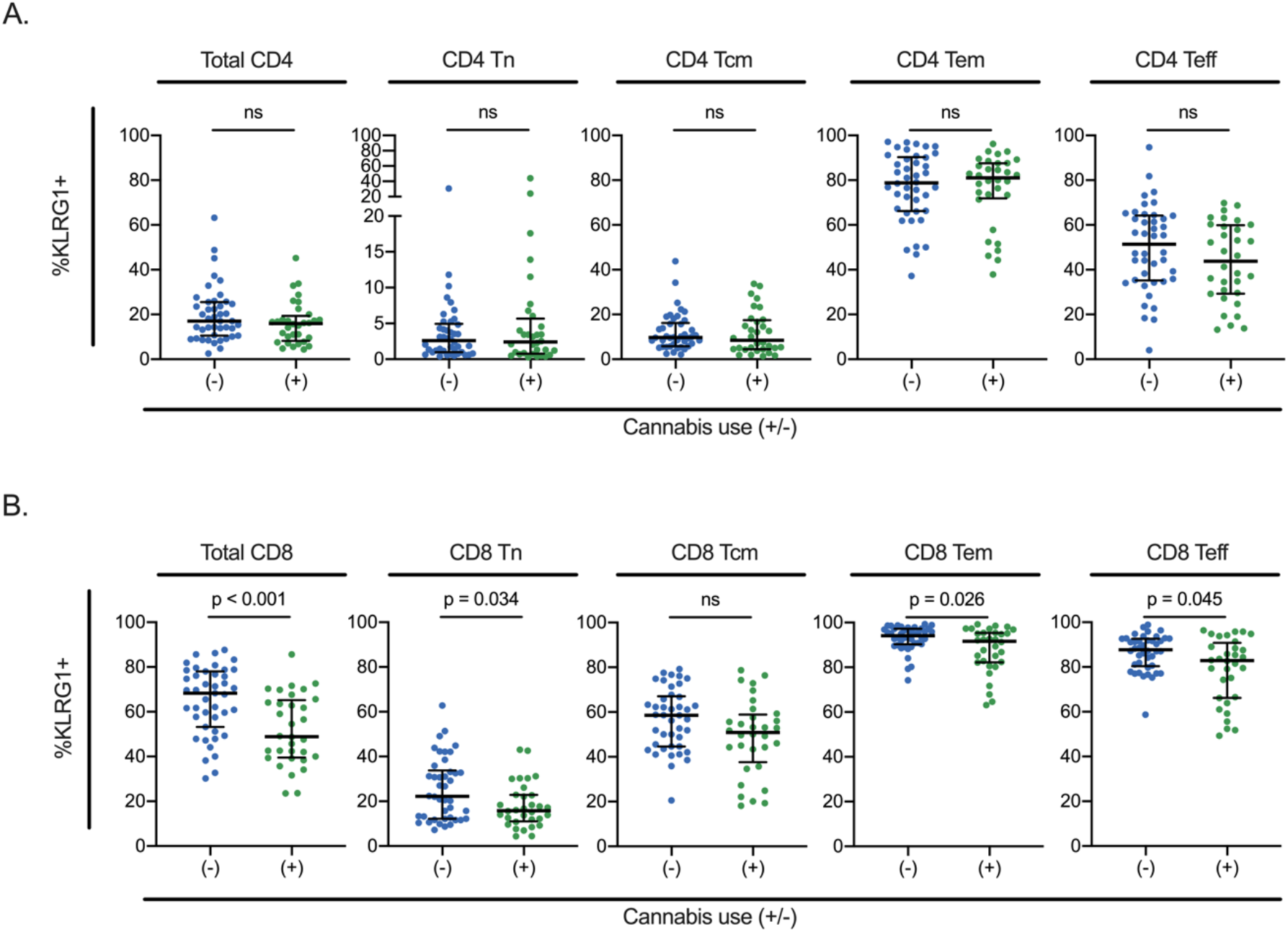
Impact of CB use on expression of the exhaustion/senescence marker KLRG1 in CD4 and CD8 T cell subsets. Expression level (percent positive) for KLRG1 is shown for total and specific subsets (Tn, Tcm, Tem, Teff) of CD4 T cells **(A)** and CD8 T cells **(B)**. Each dot represents an individual participant. Statistical significance determined using a Mann-Whitney test. P values for comparisons where CB users were significantly different (p < 0.05) to non-user controls are displayed. ns = not significant (p ≥ 0.05). Median and IQR are displayed for each dataset.

### HIV-specific T cell responses are intact in CB users

CBs have immunosuppressive and anti-inflammatory effects on the human immune system (59). To further examine the potential impact of CB use on HIV-specific functional T cell responses, we stimulated PBMCs from each study participant with a pool of HIV-derived peptides for 6 h, followed by staining for intracellular cytokines (IFN-γ, TNF-α and IL-2) as well as for the degranulation marker CD107. We then quantified the fraction of cells that were single positive for each cytokine/marker as well as the frequency of polyfunctional T cells (percent IL-2^+^/TNFα^+^/IFNγ^+^ cells within the CD4 or CD8 T cell subsets) (**Figure 6**). Within CD4 T cells from non-users, HIV peptide stimulation caused a significant increase in the fraction of positive cells for each of the cytokines as well as for CD107, indicating a robust HIV-specific T cell response. Furthermore, the percentage of polyfunctional CD4 T cells dramatically increased from median 2.8% to 38.3% after peptide stimulation. Notably, in CB using PWH, these responses were largely intact, with IL-2 and CD107 also being induced significantly by HIV-derived peptides (median 3.0% to median 33.2%). Unlike non-users, TNFα and IFNγ expression levels in CB users were not significantly higher in stimulated cells compared to unstimulated control, but the magnitude of increase after peptide stimulation was similar: for example, for IFNγ expression there was a median difference (pre versus post peptide stimulation) of 0.016% for users vs. 0.024% for non-users.

**Figure 6.**
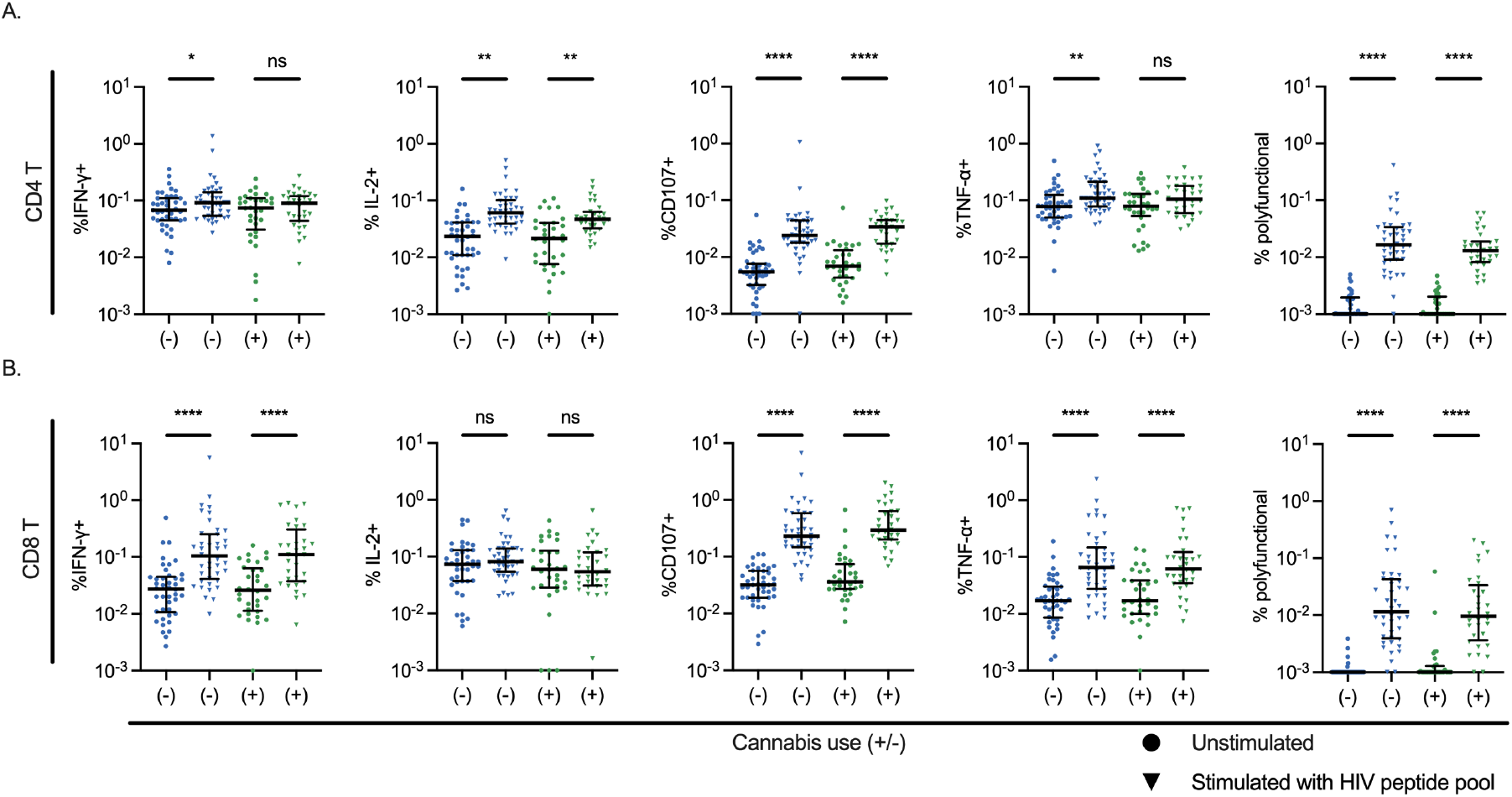
HIV-specific cytokine and effector responses in CB users are intact. PBMCs from each study participant were stimulated with a pool of HIV peptides for 6 h, then permeabilized and stained for IFNγ, TNFα, IL-2 and CD107 expression in addition to CD3, CD4 and CD8. Values represent the fraction of expressing (positive) cells within the total CD4 **(A)** and CD8 (**B)** T cell gates. Zero values or values less than 0.001 were censored at 0.001 to facilitate display on a log y axis. Polyfunctional cells are counted as the percentage of IL-2^+^/IFNγ^+^/TNFα^+^ cells within CD4 **(A)** or CD8 **(B)** T cells. Each dot represents an individual participant. Statistical significance determined using a Mann-Whitney test. P values for comparisons where CB users were significantly different (p < 0.05) to non-user controls are displayed as follows: *= p < 0.05, **= p < 0.005, ****= p <0.00005. ns = not significant (p ≥ 0.05). Median and IQR are displayed for each dataset.

We next examined HIV peptide-induced responses in CD8 T cells. In contrast with CD4 T cells, peptide stimulation did not induce significantly elevated IL-2 expression for either CB users or non-users. Nevertheless, IFNγ, TNFα and CD107 were all upregulated in stimulated CD8 T cells for both CB users and non-users, with no marked difference in the magnitude of induction across groups. After stimulation, both CB users and non-users exhibited a similar increase in the percentage of polyfunctional (IL-2^+^/TNFα^+^/IFNγ^+^) cells. Overall, these data indicate that HIV-specific CD4 and CD8 T cell responses remain intact in CB users and are not attenuated by the anti-inflammatory or immunosuppressive impact of CB use.

### Analysis of intact HIV proviral reservoir size in CB users vs non-users

We next evaluated the hypothesis that CB use impacts the frequency of the latent HIV reservoir in ART-suppressed PWH. Given their purported immunosuppressive and hypometabolic effects (42, 59–62), CBs could potentially impact reservoir size by affecting viral replication via modulation of inflammatory gene pathways that regulate HIV replication in CD4 T cells. Alternatively, CB use could affect reservoir stability during ART through modulation of immune activation and T cell maturation pathways.

Measuring the abundance of latently infected cells in CD4 T cells from ART-suppressed PWH is complicated by the presence of numerous defective proviral sequences that greatly outnumber full-length intact proviruses. We thus employed the Intact Proviral DNA Assay (IPDA) which quantifies intact HIV proviruses through a droplet digital PCR approach in which specific 5’- and 3’-proviral regions that are preserved in intact proviruses are amplified from the same vDNA molecule within a droplet, allowing intact viruses to be detected as droplets which are 5’ and 3’ double positive. This approach also enables quantitation of the frequency of 5’-defective and 3’-defective proviruses, and total HIV DNA (sum of intact, 5’-defective, and 3’-defective proviruses). To examine the impact of CB use on the frequency of the intact HIV reservoir, we performed IPDA analysis of isolated total CD4 T cells from each cohort participant. The total level of viral DNA per million CD4 T cells was not significantly different between the two cohorts, with a median frequency of 1327/10^6^ CD4 T cells for non-users and 1009/10^6^ CD4 T cells for CB users (p = 0.48, Mann-Whitney U test) (**Figure 7A**). The median level of intact (5’ and 3’ double positive) proviruses per million cells across all study participants was 50/10^6^ CD4 T cells, with a range from 5 to 1742/10^6^ CD4 T cells. Similar to total HIV DNA, median frequencies of intact proviruses were numerically less for users (48/10^6^ CD4 T cells) compared to non-users (56/10^6^ CD4 T cells), and not statistically different (p = 0.208, Mann-Whitney U test) (**Figure 7B)**. No clear differences in the percentage of intact proviruses were observed across CB users versus non-users (**Figure 7C)**. These data suggest that across the whole cohort, CB use has little apparent impact on the size of the intact HIV reservoir.

**Figure 7.**
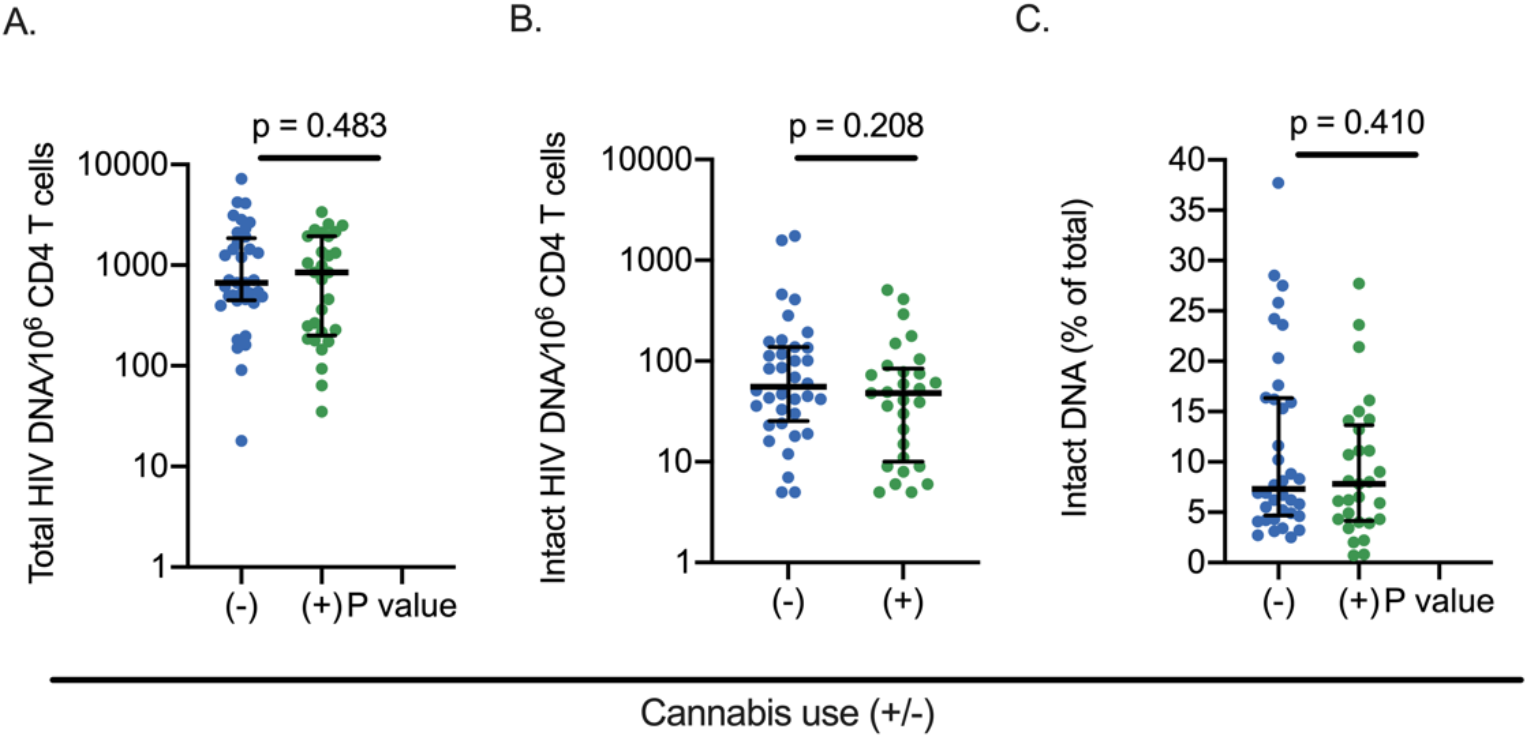
Analysis of intact HIV proviral reservoir size in CB users vs non-users. Intact Proviral DNA Assay was carried out on purified CD4 T cells from each subject to quantify **(A)** total viral DNA per million CD4 T cells (total HIV DNA/10^6^ CD4 T cells), **(B)** intact proviruses per million CD4 T cells (intact HIV DNA/10^6^ CD4 T cells), and **(C)** percent intact proviruses of total proviruses detected by IPDA for each study participant (intact DNA % of total). Each dot represents an individual participant. Statistical significance determined using a Mann-Whitney test. P values for comparisons where CB users were significantly different (p < 0.05) to non-user controls are displayed. ns = not significant (p ≥ 0.05). Median and IQR are displayed for each dataset.

We next conducted a hypothesis-generating exploratory analysis using the expression levels of various immune markers derived from our flow cytometry data to stratify cohort participants into different sub-groups and to examine whether CB use impacted the intact reservoir size within any of these groups. We focused on groups with relatively balanced numbers of participants from both the CB user and non-user cohorts. For each immune marker or demographic characteristic, we considered all possible thresholds values that lead to cannabis user and non-user groups size of at least 15 samples (note that for these experiments our cohort size is 66 rather than 75 due to nine samples failing to produce an IPDA value). Among these thresholds for each marker, we chose one that maximized the median difference between cannabis user and non-user cohorts. From this approach, we identified four subgroups of PWH within which CB users exhibited a statistically significantly smaller (p < 0.05) intact reservoir size (**Figure 8, Table 2**). Interestingly, three of these groups were identified based on the expression of surface markers in NK cells. These groups were: 1) participants with a low frequency of CCR7^+^/IFNγ^-^/IL-2^-^/TNFα^+^ CD4 T cells (**Figure 8A**), 2) participants with a low frequency of NKG2A^-^/CD16^+^ cells in the NK cell subset (**Figure 8B**), 3) participants with a high frequency of NKG2A^+^/CD16^+^ cells in the NK cell subset (**Figure 8C**), and 4) participants with a high frequency of NKG2A^+^ cells in the NK cell subset (**Figure 8D**). Overall, these data indicate that, although the impact of CB use on reservoir size was not statistically significant across the entire cohort, there were sub-groups of CB users for whom CB use may be associated with reduced intact provirus reservoir size. One intriguing hypothesis based on this exploratory analysis is that the NK cell compartment may play a role in regulating the impact of CB on the size of the HIV reservoir. This hypothesis will require further investigation and validation in additional cohorts.

**Table 2:** Subgroups of participants with different intact reservoir size for CB users.

| Split | Number of samples | Number of non CB users | Number of CB users | Non CB users median Intact vDNA/10<sup>6</sup> | CB users median Intact vDNA/10<sup>6</sup> | Absolute value of median difference | Mann Whitney U Test Statistic | Mann Whitney U Test p value |
| --- | --- | --- | --- | --- | --- | --- | --- | --- |
| NK NKG2A+ freq > 28.1% | 33 | 18 | 15 | 109 | 39 | 70 | 60.5 | 0.006 |
| NK NKG2A-/CD16+ freq ≤ 55.9% | 32 | 16 | 16 | 100.5 | 40 | 60.5 | 69 | 0.025 |
| NK NKG2A+/CD16+ freq > 15.4% | 32 | 16 | 16 | 93 | 37.5 | 55.5 | 67.5 | 0.021 |
| CD4 T CCR7+/IFNγ-/IL2-/TNFα+ freq ≤ 0.000645% | 45 | 24 | 21 | 106.5 | 41 | 65.5 | 161.5 | 0.039 |

IPDA data for four subsets of study participants within which CB users exhibit smaller intact HIV reservoir size than non-users are shown. Subsets were selected based on the frequency of expression of immune markers within certain immune populations (shown in left column).

**Figure 8.**
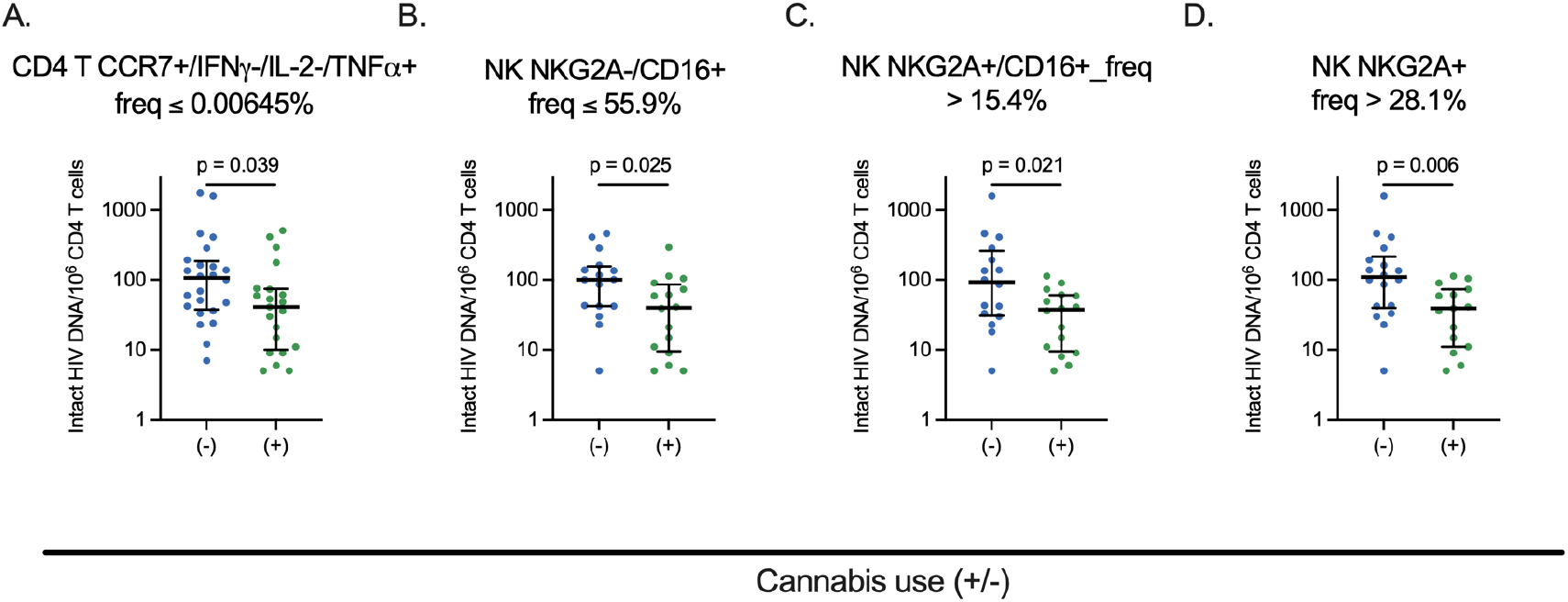
Analysis of intact HIV proviral reservoir size in CB users vs non-users within specific subgroups. Intact Proviral DNA Assay was carried out on purified CD4 T cells from each subject to quantify intact proviruses per million CD4 T cells (intact HIV DNA/10^6^ CD4 T cells) for each study participant. Data are shown from specific subsets of study participants based on flow cytometry expression patterns described above each plot (**A-D)**. Each threshold was selected to generate groups with at least 15 participants from each cohort and to maximize the difference in median intact proviruses per million cells. Each dot represents an individual participant. Statistical significance determined using a Mann-Whitney test. p values for comparisons where CB users were significantly different (p < 0.05) to non-user controls are displayed. ns = not significant (p ≥ 0.05). Median and IQR are displayed for each dataset.

### Data visualization and machine learning analysis of immune marker expression and reservoir size for CB users

Next, we used data visualization and machine learning tools to analyze the overall patterns of surface marker expression and IPDA data to determine whether this data could be used to separate CB users as distinct from non-users. We reasoned that such an approach might provide important insights into differences between the cohorts that may not be apparent from more traditional analyses. For the visualization, our data consists of 230 total features, among which 216 features are derived from the flow cytometry data, 13 from demographic data, and 1 from IPDA intact provirus data. Categorical features, such as race were converted to binary variables.

First, the abilities of individual surface markers to classify individuals as being CB users or non-users were evaluated using Receiver Operating Characteristic (ROC) curves and the Area Under the Curves (AUC) metric. To create ROC curves, we plotted the true positive rate (TPR) against the false positive rate (FPR) at different possible threshold settings for each individual surface marker frequency of expression. TPR (or sensitivity) measures the proportion of positive samples that are correctly identified, while FPR measures the proportion of negative samples that are incorrectly identified as positive. A perfect model is represented by a plot that begins vertically then moves to the right at the top of the plot, meaning that there are no false negatives and no false positives, while a random guessing model corresponds to a diagonal line on the ROC curve. We also computed Area Under the Curve (AUC), which is the probability that a model will be able to distinguish between a randomly chosen positive sample and a randomly chosen negative sample. AUC is a summary statistic of ROC curves, and a perfect model has an AUC of 1, while a random guess (diagonal line) has an AUC of 0.5. For our experiments, we plotted ROC curves for each feature (frequency of expression for given surface marker or combination of markers in a population of cells) in our data separately (**Figure 9A**). We consider a feature to be able to discriminate between cannabis and non-cannabis users if its AUC value is greater or equal to 0.6. In total, we found 46 features among 230 that had an AUC value greater or equal to 0.6 (**Figure 9A, Table 3**). We show four features with the highest AUC value in color in **Figure 9A**. Overall, KLRG1, and PD-1 expression were prominently represented among the sets of features that best distinguished cannabis users from non-users, highlighting the impact of cannabis use on T cell exhaustion and senescence. The top two features were both derived from CD8 T cells and were 1) percent KLRG1^-^/PD-1^-^ cells within CD8 T cells (AUC = 0.723), and 2) percent PD-1^-^/CCR7^+^ cells within CD8 T cells (AUC = 0.722). These findings indicate that these methods can be used to identify host features that best distinguish different cohorts, and that KLRG1, PD-1 and CCR7 expression are key features that differentiate CB users from non-users.

**Table 3:** Area under the curve values of ROC curves for individual features

| Feature | AUC | Feature | AUC | Feature | AUC |
| --- | --- | --- | --- | --- | --- |
| CD8 T KLRG1-/PD1- freq | 0.7229 | CD8 T HLA-DR+ freq | 0.6616 | CD8 T CM CD38-/HLA-DR- freq | 0.6230 |
| CD8 T PD1-/CCR7+ freq | 0.7215 | CD8 T E freq | 0.6613 | CD4 T KLRG1+/CD27- freq | 0.6212 |
| CD8 T KLRG1+ freq | 0.7096 | CD4 T CM CD38+/HLA-DR- freq | 0.6613 | CD8 T EM CD38-/HLA-DR- freq | 0.6194 |
| CD8 T KLRG1-/CD27+ freq | 0.7085 | CD4 T KLRG1-/PD1- freq | 0.6522 | CD8 T CD38+/HLA-DR+ freq | 0.6158 |
| CD8 T CM PD1+ freq | 0.6970 | CD4 T EM PD1+ freq | 0.6497 | CD4 T E CD38-/HLA-DR- freq | 0.6158 |
| CD8 T KLRG1+/CD27- freq | 0.6959 | CD8 T KLRG1+/PD1- freq | 0.6483 | CD4 T EM freq | 0.6144 |
| CD4 T N CD38-/HLA-DR+ freq | 0.6926 | CD8 T N CD38-/HLA-DR- freq | 0.6483 | CD4 T KLRG1+/PD1+ freq | 0.6129 |
| CD4 T CM CD38-/HLA-DR+ freq | 0.6919 | CD8 T CD38-_/HLA-DR+ freq | 0.6403 | CD8 T KLRG1+/PD1+ freq | 0.6111 |
| CD8 T CD27+ freq | 0.6797 | CD8 T N PD1+ freq | 0.6385 | CD8 T polyfxn freq | 0.6104 |
| CD4 T N CD38+/HLA-DR- freq | 0.6789 | CD4 T CD27+ freq | 0.6367 | CD4 T N freq | 0.6093 |
| CD8 T N freq | 0.6717 | CD4 T PD1+/CCR7- freq | 0.6338 | CD8 T N CD38+/HLA-DR+ freq | 0.6050 |
| CD4 T PD1+ freq | 0.6674 | CD8 T E CD38-/HLA-DR- freq | 0.6320 | CD4 T KLRG1-/CD27+ freq | 0.6035 |
| CD4 T N CD38-/HLA-DR- freq | 0.6656 | CD4 T CM PD1+ freq | 0.6302 |  |  |
| CD4 T PD1-/CCR7+ freq | 0.6616 | CD8 T PD1-/CCR7- freq | 0.6291 |  |  |

ROC curves were used to determine the ability of individual features of the flow cytometry dataset to accurately classify CB users vs non-users. AUC=area under the ROC curve. N=naïve, CM=central memory, E=effector, polyfxn= polyfunctional (IL-2+/TNFα +/IFNγ + cells).

**Figure 9.**
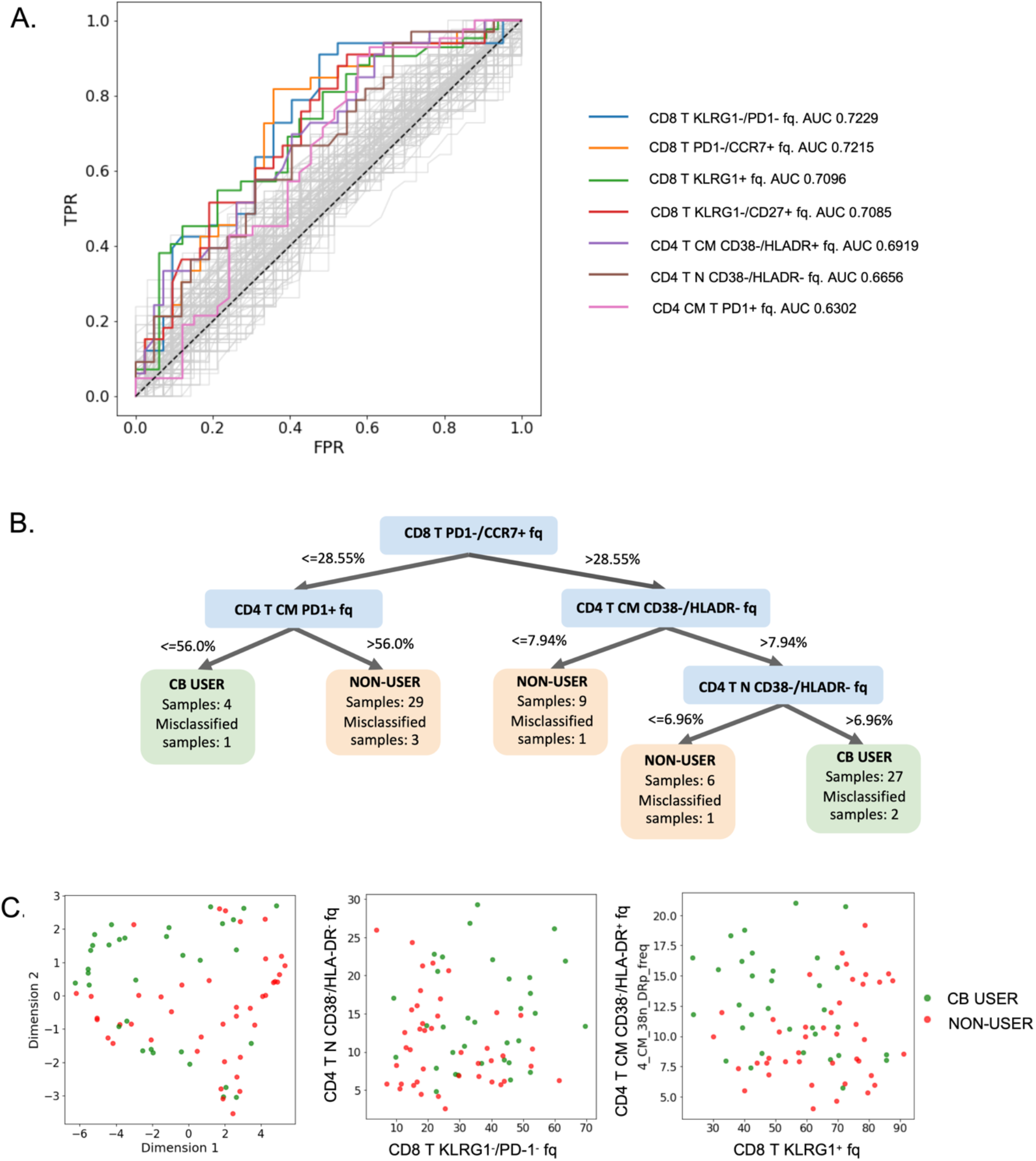
Machine learning analysis of IPDA and flow cytometry data to predict cannabis use. **A**. ROC curves and corresponding AUC values for 230 features derived from demographics, IPDA and flow cytometry analysis of the cohort (frequency of cell subsets based on surface marker expression). Axes represent the true positive rate (TPR) and the false positive rate (FPR) for each feature-derived model for classifying study participants as CB users or non-users. Striped black line corresponds to the ROC curve of an uninformative model. In color, we depict ROC curves of features with highest AUC values and as well as features used in Decision Tree visualization of the dataset. **B**. Visualization of the dataset using the Decision Tree algorithm GOSDT (63). This tree correctly describes 89.33% of the dataset. **C**. Visualization of the dataset based on application of PaCMAP dimension reduction (left panel) and scatter plot of two sets features with high AUC values: the frequency of CD38^-^/HLA-DR^-^ cells within the CD4 Tn population versus the frequency of KLRG1^-^/PD-1^-^ cells within the total CD8 T cell gate (middle panel), the frequency of CD38^-^/HLA-DR^+^ cells within the CD4 Tcm population versus the frequency of KLRG1^+^ cells within the CD8 T cell population (right panel). N=Naïve, CM= Central memory.

As an additional analysis, we employed a decision tree method to visualize the data cohort and observe differences between CB users and non-users with simple rules. Specifically, we used Generalized and Scalable Optimal Sparse Decision Trees (GOSDT) (63) to identify a tree that could optimally classify CB users versus non-users using the demographics and flow cytometry data from each subject. GOSDT produces an optimal decision tree over all possible trees given the user-defined parameters, such as regularization that controls sparsity or decision tree depth (**Figure 9B**). This approach generated a tree that correctly described approximately 90% of the dataset, with only 8 out of 75 samples misclassified. This algorithm created five “leaves”, meaning that it places the data into five groups, classifying members of each group the same way. We use two criteria for hyperparameter tuning: high percentage of correctly described samples and absence of leaves with one or two samples in them. The resulting tree has a maximum of three splits and has leaf support (number of samples in each leaf) higher than 5% of the overall cohort. Notably, these decision splits were determined by a specific set of flow cytometry markers – in particular, PD-1/CCR7 expression in CD8 T cells and PD-1/CD38/HLA-DR expression in central memory CD4 T cells played prominent roles in the decision tree visualization (**Figure 9B**).

Finally, we examined the ability of combinations of features to distinguish CB users with dimension reduction tools. Dimension reduction techniques have been used to visualize data for many real-world datasets. In our analyses, we performed well-known dimension reduction techniques such as t-SNE (64), UMAP (65), and PaCMAP (66). Among all of these, we found that PaCMAP visualization produced the most distinct clusters, since this algorithm was built to preserve both global and local structure of the dataset (**Figure 9C, left panel, Figure S1**). CB users and non-users exhibited spatially distinct areas of the plots, indicating differences between the cohorts, although some overlap was also visible. We also examined whether visualizing binary combinations of features from the dataset with high AUC values could allow visual separation of the cohorts on a scatter plot **(Figures 9C, middle and right panels)**. We observed that visualization of the following combinations of features demonstrated separation of CB users and non-users: 1) percent KLRG1^-^/PD-1^-^ cells within CD8 T vs percent CD38^-^/HLA-DR^-^ within CD4 Tn cells, and 2) percent KLRG1^+^ within CD8 T cells vs percent CD38^-^/HLA-DR^+^ within CD4 Tcm cells allowed us to observe separate clusters of CB users and non-users.

In conclusion, using data science approaches, such as ROC curves and AUC, decision trees, and dimension reduction techniques for visualization, we were able to find ways to visualize and highlight key markers that distinguish CB users from non-users. More specifically, we found that the frequency of specific subsets of cells (CD8 T KLRG1^-^/PD-1^-^, CD8 T PD-1^-^/CCR7^+^, CD4 Tn CD38^-^/HLA-DR^-^, CD4 Tcm CD38^-^ /HLA-DR^+^, CD8 T KLRG1^+^) can be used to discriminate CB users from non-users. Methods we used here may be generally useful to understand the complex immunological outcomes of a given drug exposure.

## Discussion

Despite the effectiveness of antiretroviral therapy, a reservoir of latently infected cells persists in treated individuals and precludes HIV cure (67). Furthermore, ART-treated PWH maintain persistently high levels of chronic inflammation, immune activation and exhaustion that likely contribute to accelerated immune aging, increased morbidity and reduced life expectancy in these individuals (10, 58, 68). The driving forces behind this residual immunological dysfunction remain unresolved but could be at least partially driven by residual viral transcription and protein expression promoting activation of innate, or adaptive immune signaling pathways. If this hypothesis is correct, the latent reservoir itself could be a key driver of persistent immune activation in PWH. Additionally, increased immune activation may drive clonal expansion of latently infected cells, thereby promoting maintenance of the reservoir. The bi-directional viral/immune dynamics of the HIV reservoir will need to be fully clarified by additional studies. However, it is possible that strategies to limit residual immune activation could limit maintenance of the reservoir through clonal expansion, or conversely, that elimination of the viral reservoir could mitigate immune activation.

Despite the high prevalence of cannabis use in PWH, there are only a handful of studies to date that have examined the impact of cannabis use on the immune system or on the viral reservoir in PWH (20, 35, 38, 40, 48, 50, 69, 70). Several of these prior studies have indicated that CB impacts the immune systems of PWH. For instance, PWH who use CB have a lower frequency of CD16^+^ inflammatory monocytes and plasma IP10 compared to non-users (48). Manuzak et al. examined the expression of activation markers in total CD4 and CD8 T cells, and observed that heavy CB users, but not moderate users, exhibited a reduced frequency of activated (HLA-DR^+^/CD38^+^) cells for both CD8 and CD4 T cells (35). This study also observed reduced abundance of CD11c^+^/CD123^-^ classical dendritic cells (cDCs) in moderate and heavy users, as well as lower cytokine (IL-23, IL-6, IL-10 TNFα) production in antigen-presenting cells (APCs) (CD3^-^/CD20^-^/HLA-DR^+^) in heavy users, but no change to CD123^+^ plasmacytoid DCs (pDCs). Animal studies have also indicated a possible role of CBs in reducing inflammation during infection – THC administration has been shown to reduce intestinal inflammation in SIV infected macaques (49). However, some studies have contradicted these findings and have observed either no impact of CB use on inflammation in PWH, or increased levels of some inflammatory markers (39, 71).

Our findings here are broadly consistent with the notion that CB use is associated with reduced immune activation in CB using PWH and also provide additional important insights. Consistent with previous reports, we observed no change in the proportion of total CD4 and CD8 T cells in peripheral blood (40). However, we observed an elevated proportion of naïve CD8 T cells and reduced effector CD8 T cells in CB users. Furthermore, there was significantly reduced expression of the T cell senescence marker KLRG1 in total CD8 T cells, and in most CD8 T cell memory subsets of CB users compared to non-users. Overall, these data strongly support the notion that CB use is associated with reduced T cell immune activation and senescence in ART-suppressed PWH. Cannabinoids may thus provide some immunological benefit to PWH and could potentially help to prevent non-AIDS comorbidities. Cannabinoids should be further explored as a potential therapeutic for PWH.

In addition to high dimensional immunophenotyping, this study also provides a novel assessment of the impact of CB use on HIV-specific T cell responses. We assessed HIV antigen-induced *ex vivo* expression of TNFα, IL-2, IFNγ, as well as surface levels of the degranulation marker CD107. Overall, we observed that these responses are preserved in the CB-users relative to non-users in our study. It remains possible that CB use could cause transient reduction in adaptive responses to HIV during CB use *in vivo*, or impact adaptive responses in a manner that is not captured using *ex vivo* assays. This observation suggests that T cells from PWH who use cannabis may still retain functionality for potential killing and clearance of residual infected cells that express antigen.

The mechanism by which CB use reduces T cell activation and exhaustion in PWH is unclear. Some *in vitro* studies show that CBs can limit T cell activation and proliferation, indicating a potential direct effect on T cells (30, 59, 72). The cannabinoid receptor CB_2_ is expressed by T cells and is associated with modulation of several signaling pathways related to activation of cellular proliferation and metabolism (73). The relationship between the canonical CB-receptor driven proliferative cellular signaling and data suggesting that CBs limit immune activation and T cell proliferation deserves further study. CB_2_ is also abundantly expressed in numerous other immune cell types which may influence HIV specific T cells indirectly. For example, CB use has been shown to impact the abundance of monocyte and DC subsets as well as cytokine secretion by antigen presenting cells in PWH (35, 48). THC can also inhibit activation of T cells through suppression of innate antiviral cytokines such as type 1 interferon (74, 75).

Interestingly, a previous study that examined longitudinal abundance of HIV vDNA in PWH observed a significantly more rapid viral DNA (vDNA) decay in CB users after ART initiation (70). This trend was not observed when considering PWH that use additional drugs of abuse (DOA), suggesting that positive benefits of CB use could be counteracted by other DOA. Consistent with this notion, tobacco exposure seems to reverse anti-inflammatory effects of CB use (50). These observations suggest that additional DOA use could be a significant confounder for understanding the impact of CB use.

Our study also demonstrates a potential impact of CB use on the frequency of the intact latent HIV reservoir. Specifically, CB-users had numerically lower frequencies of total and intact HIV DNA in CD4 T cells, although these differences were not statistically significant. In an exploratory hypothesis-generating analysis, we found that CB use may have a more significant impact on specific subsets of PWH. In particular, there was a more significant association of CB use with lower intact HIV DNA frequencies in when only study participants with a high frequency of NKG2A^+^ NK cells were examined (p = 0.006).

Additional studies in larger cohorts will be needed to confirm whether CB use has an impact on the size of the intact latent HIV reservoir. One possible way in which CB use could limit the size of the HIV reservoir is by inhibiting HIV replication before ART, leading to fewer total infected cells. Consistent with this hypothesis, some evidence indicates that high CB use is associated with reduced HIV RNA levels in in newly infected people (69). Additionally, THC dosing of SIV-infected macaques leads to lower viral loads and better survival, although this effect was only observed for male animals (41). Also, HIV replication in macrophages can be directly inhibited by CB_2_ agonists (31). CB could also impact the HIV reservoir size post-ART through immunological mechanisms. For example, CB could limit activation of inflammatory pathways that drive clonal expansion of latently infected cells. Alternatively, by limiting immune exhaustion and senescence, CB could promote increased HIV-specific immune functionality, leading to more rapid immune-dependent elimination of infected cells from the reservoir. Our observation that study subjects with high frequency of expression of NKG2A in the NK cell compartment exhibits a large magnitude of impact of CB on HIV reservoir size is intriguing. NK cells are known to be able to kill HIV infected cells (76, 77), and these cells are also affected by exposure to cannabinoids (78). Additional work will be needed to clarify the relationship between NK cells, CB use and the HIV reservoir.

Our study should also be considered in the context of some limitations. The size of our CB-using and control cohorts may be too small to observe more subtle effects on immune populations and on the HIV reservoir. Furthermore, our study does not distinguish between heavy CB users versus moderate users, which could obscure more significant effects observed in heavy users. Nevertheless, our study provides evidence that CB can have a significant impact on the immune status of PWH and may be associated with reduced size of latent HIV reservoir in some PWH, although further study is needed. An important open question regarding the impact of CB on the HIV reservoir and on the immune system of PWH is the identification of specific CB constituents that mediate these outcomes. Cannabis contains several CBs, including– THC (the main psychoactive ingredient), cannabidiol (CBD), and minor cannabinoids such as cannabigerolic acid (CBGA). Animal model experiments will likely be necessary to establish which CBs could provide the most immune benefit to PWH, and to more fully assess the benefits and risks of CB use on non-immune tissues and physiological and psychological processes.

## Methods

### Study Approval

This study was reviewed and approved by the IRBs from both Duke University and UNC (00107328).

### Peptide Stimulation

Peripheral whole blood was obtained from IRB-approved donors using ACD vacutainer tubes (BD Biosciences), and peripheral blood mononuclear cells (PBMCs) were isolated using Ficoll density centrifugation (GE HealthCare). PBMCs were counted and viably cryopreserved in liquid nitrogen vapor (10% DMSO, 90% heat-inactivated FBS). Thawed PBMCs were rested for 6 h prior to stimulation in R10, which is RPMI-1640 media containing 10% heat-inactivated (HI)-FBS (Gibco) and 1x penicillin-streptomycin-glutamine (Gibco), at 37°C and 5% CO_2_. Cells were stimulated with HIV Gag, Pol, and Env PepMixes (JPT Laboratories) at 0.2 ug/mL final concentration of each in R10 at a cell concentration of 2 × 10^7^ cells/mL for 6 h. The final concentration of DMSO was less than 0.2%. Cells were stimulated in the presence of brefeldin A (BFA) and monensin per manufacturer protocol (BD Bioscience). CD107a-PE (H4A3 clone, BL) was included during stimulation.

### Flow Cytometry

Cell preparation and staining followed methods previously described (79). All monoclonal antibodies were titrated to optimal signal-to-noise ratio on PBMCs prior to use, assuming a 50μL staining volume. Cell viability was determined using Zombie-NIR Fixable Viability Dye (0.4 μL per 50 μL staining volume; Biolegend). Antibodies used for surface staining were as follows: KLRG1-BV421 (SA231A2 clone, Biolegend, CD45RA-PacBlue (H100 clone, BL), CD8-BV570 (RPA-T8 clone, Biolegend), CD127-BV605 (A019D5 clone, Biolegend), CD56-BV650 (5.1H11 clone, Biolegend), CCR7-BV711 (G043H7 clone, Biolegend), CD27-BV750 (O323 clone, Biolegend), PD1-VioBright515 (REA165 clone, Miltenyi), NKG2A-PE-Vio615 (REA110 clone, Miltenyi), CD16-PerCP-Cy5.5 (33G8 clone, Biolegend), CD38-PCPeF710HB7 clone, TF), CD14-SparkNIR685 (63D3 clone, Biolegend), CD19-SparkNIR685 (HIB19 clone, Biolegend), HLA-DR-APC-F750 (L243 clone, Biolegend). Antibodies used for intracellular staining were as follows: CD3-BV480 (UCHT1 clone, BD), CD4-PerCP (L200 clone, BD), IFN-g-PE-Cy7 (4S.B3 clone, Biolegend), IL-2-APC (MQ1-17H12 clone, Biolegend), TNFa-AF700 (Mab11 clone, Biolegend). Sample acquisition was performed on a Cytek Aurora spectral flow cytometer. Flow cytometry analysis was performed in FlowJo v10.8 software.

### Intact Proviral DNA analysis (IPDA)

Cryopreserved samples of PBMCs from each study participant were viably thawed. A portion was dedicated to immunophenotyping and functional assays described above, and the remainder were subjected to total CD4 T cell negative selection with the StemCell Technologies EasySep™ Human CD4+ T Cell Enrichment Kit (Cat# 19052). A median of 4.4 million CD4 T cells (Q1 3.5 million, Q3 5.1 million) were isolated for each participant with a median lymphocyte purity of 96.5% (Q1 95%, Q3 98%) measured with a Sysmex hematology analyzer. CD4 T cell DNA was extracted using the QIAamp DNA Mini Kit and quantified on a NanoDrop 1000 (Thermo Fisher Scientific). IPDA was performed as originally described (45), with a validated PCR annealing temperature modification to increase signal to noise ratio (80). A detailed protocol for the assay is described in detail elsewhere (45, 80). Gating for positive droplets was set using negative (DNA elution buffer and HIV-seronegative CD4 T-cell DNA), and positive (Integrated DNA Technologies gblock amplicon) positive control wells processed in parallel. DNA shearing index values were similar to those reported previously (median, 0.33; interquartile range [IQR], 0.31–0.34). A median of 1.1 × 10^6^ (IQR, 9.4 × 10^5^–1.2 × 10^**6**^) CD4 T-cell equivalents (2 RPP30 copies = 1 CD4 T cell equivalent) were evaluated per sample. Samples for which either the PS, RRE, or both probes failed to amplify were excluded from analysis (9 of 75 participants evaluated, or 12%). Proviral frequencies less than 5 copies/million CD4 T cells were left-censored as previously described (80).

### Statistical analysis and machine learning

The two cohorts consisted of 33 CB-using and 42 non-using people with HIV (PWH). Statistical comparisons between the cohorts for flow cytometry and IPDA analysis were carried out using a Mann-Whitney U test with the threshold of significance as p < 0.05. The primary endpoint for the IPDA analysis was the frequency of intact proviruses across the cohorts, with a secondary exploratory analysis of selected subgroups based on different thresholds of frequency for immune cell populations. Given that the secondary analysis was exploratory and hypothesis-generating, no correction for multiple comparisons was performed.

ROC curves for each cytometry marker and demographics feature were created by plotting true positive rate (TPR) versus false positive rate (FPR) for every threshold value (frequency of expression). We further computed AUC using the trapezoidal rule given TPR and FPR arrays. For Decision Tree visualization we used the GOSDT algorithm to construct a tree that could identify patterns in dataset and visualize cannabis using subjects versus non-users (63). In the GOSDT setting, we set the regularization parameter to 0.0134, the depth budget to 4, and time limit to 5000. The hyperparameters were chosen to ensure that the resulting tree describes the data with high accuracy and each leave has at least three samples. Lastly, we utilized dimension reduction tools such as t-SNE (64), UMAP (65), and PaCMAP (66) to visualize the dataset. We standardized the data before dimension reduction, set the random state to sixteen for reproducibility, and used PCA for initialization of lower dimensions. We set the number of neighbors to six for PaCMAP, early exaggeration to eight and perplexity to 40 for t-SNE, number of neighbors to four and minimal distance to 0.2 for UMAP.

## Author contributions

Designed the study – EPB, DM Murdoch, Conducted experiments – SDF, ADCV, AM. Analyzed data – EPB, DM Murdoch, AR, EW, LS, CDR, SDF. Wrote manuscript: EPB, SDF, NA, LS, DM Murdoch, DM Margolis.

## Acknowledgements

This work was supported by the following grants from the National Institutes of Health: NIAID R01 AI143381 (EPB), NIAID UM1 AI164567 (DM Margolis), NIDA R61 DA047023 (EPB), NIDA R01 DA054994 (CDR), NIAID F30 AI145588 (SDF).

---

**Figure S1:**
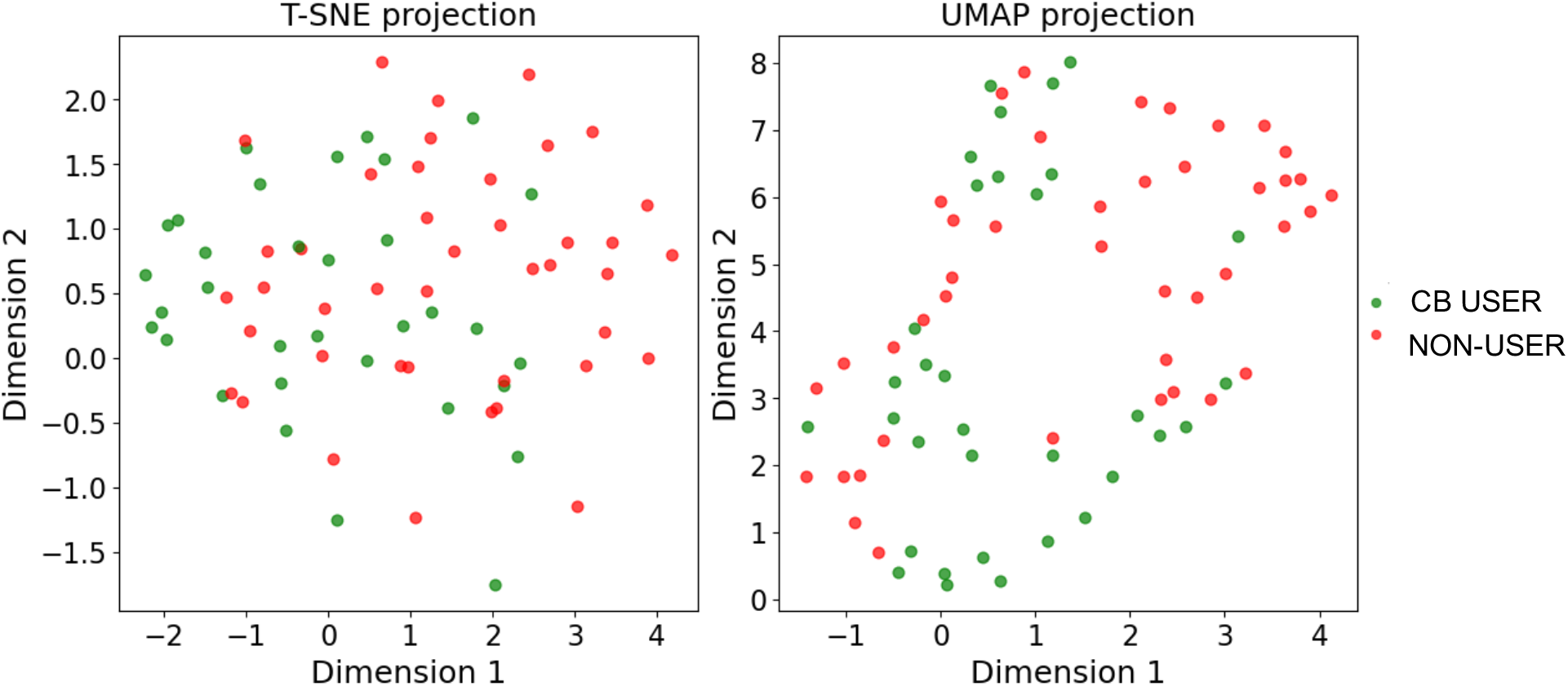
The overall dataset is displayed using t-SNE (left panel) and UMAP (right panel). Each dot represents a study participant. CB users indicated by green dots, non-users by red dots.

## References

1. Eisinger RW, Fauci AS. Ending the HIV/AIDS Pandemic. Emerg Infect Dis 2018;24(3):413–416.

2. Chun T-W et al. Presence of an inducible HIV-1 latent reservoir during highly active antiretroviral therapy. PNAS 1997;94(24):13193–13197.

3. Chun T-W et al. Early establishment of a pool of latently infected, resting CD4+ T cells during primary HIV-1 infection. Proc Natl Acad Sci U S A 1998;95(15):8869– 8873.

4. Finzi D et al. Identification of a Reservoir for HIV-1 in Patients on Highly Active Antiretroviral Therapy. Science 1997;278(5341):1295–1300.

5. Siliciano JD et al. Long-term follow-up studies confirm the stability of the latent reservoir for HIV-1 in resting CD4+ T cells. Nat Med 2003;9(6):727–728.

6. N S-S et al. Quantitation of replication-competent HIV-1 in populations of resting CD4+ T cells. Journal of virology 2014;88(24). doi:10.1128/JVI.01900-14

7. Wang Z et al. Expanded cellular clones carrying replication-competent HIV-1 persist, wax, and wane. Proc Natl Acad Sci U S A 2018;115(11):E2575–E2584.

8. Cohn LB, Chomont N, Deeks SG. The Biology of the HIV-1 Latent Reservoir and Implications for Cure Strategies. Cell Host & Microbe 2020;27(4):519–530.

9. Papagno L et al. Immune activation and CD8+ T-cell differentiation towards senescence in HIV-1 infection. PLoS Biol 2004;2(2):E20.

10. Trautmann L et al. Upregulation of PD-1 expression on HIV-specific CD8+ T cells leads to reversible immune dysfunction. Nat Med 2006;12(10):1198–1202.

11. Bofill M et al. Increased numbers of primed activated CD8+CD38+CD45RO+ T cells predict the decline of CD4+ T cells in HIV-1-infected patients. AIDS 1996;10(8):827–834.

12. Utay NS, Hunt PW. Role of immune activation in progression to AIDS. Curr Opin HIV AIDS 2016;11(2):131–137.

13. Nasi M et al. Ageing and inflammation in patients with HIV infection. Clin Exp Immunol 2017;187(1):44–52.

14. Almeida C-AM, Price P, French MAH. Immune activation in patients infected with HIV type 1 and maintaining suppression of viral replication by highly active antiretroviral therapy. AIDS Res Hum Retroviruses 2002;18(18):1351–1355.

15. Wada NI et al. The effect of HAART-induced HIV suppression on circulating markers of inflammation and immune activation. AIDS 2015;29(4):463–471.

16. Marcus JL et al. Comparison of Overall and Comorbidity-Free Life Expectancy Between Insured Adults With and Without HIV Infection, 2000-2016. JAMA Netw Open 2020;3(6):e207954.

17. Hunt PW et al. Gut epithelial barrier dysfunction and innate immune activation predict mortality in treated HIV infection. J Infect Dis 2014;210(8):1228–1238.

18. Han BH, Palamar JJ. Marijuana use by middle-aged and older adults in the United States, 2015-2016. Drug Alcohol Depend 2018;191:374–381.

19. Nichols SL et al. Concordance between self-reported substance use and toxicology among HIV-infected and uninfected at risk youth. Drug Alcohol Depend 2014;134:376–382.

20. Lutge EE, Gray A, Siegfried N. The medical use of cannabis for reducing morbidity and mortality in patients with HIV/AIDS. Cochrane Database Syst Rev 2013;(4):CD005175.

21. Galiègue S et al. Expression of central and peripheral cannabinoid receptors in human immune tissues and leukocyte subpopulations. Eur. J. Biochem. 1995;232(1):54–61.

22. Massi P, Vaccani A, Parolaro D. Cannabinoids, immune system and cytokine network. Curr. Pharm. Des. 2006;12(24):3135–3146.

23. Munro S, Thomas KL, Abu-Shaar M. Molecular characterization of a peripheral receptor for cannabinoids. Nature 1993;365(6441):61–65.

24. Pini A et al. The role of cannabinoids in inflammatory modulation of allergic respiratory disorders, inflammatory pain and ischemic stroke. Curr Drug Targets 2012;13(7):984–993.

25. Börner C, Smida M, Höllt V, Schraven B, Kraus J. Cannabinoid receptor type 1- and 2-mediated increase in cyclic AMP inhibits T cell receptor-triggered signaling. J. Biol. Chem. 2009;284(51):35450–35460.

26. Ye L, Cao Z, Wang W, Zhou N. New Insights in Cannabinoid Receptor Structure and Signaling. Curr Mol Pharmacol 2019;12(3):239–248.

27. Porcella A, Gessa GL, Pani L. Delta9-tetrahydrocannabinol increases sequence-specific AP-1 DNA-binding activity and Fos-related antigens in the rat brain. Eur. J. Neurosci. 1998;10(5):1743–1751.

28. Hu Y et al. Single-cell Transcriptome Mapping Identifies Common and Cell-type Specific Genes Affected by Acute Delta9-tetrahydrocannabinol in Humans. Sci Rep 2020;10(1):3450.

29. Ehrhart J et al. Stimulation of cannabinoid receptor 2 (CB2) suppresses microglial activation. J Neuroinflammation 2005;2:29.

30. Yuan M et al. Delta 9-Tetrahydrocannabinol regulates Th1/Th2 cytokine balance in activated human T cells. J. Neuroimmunol. 2002;133(1–2):124–131.

31. Ramirez SH et al. Attenuation of HIV-1 replication in macrophages by cannabinoid receptor 2 agonists. J. Leukoc. Biol. 2013;93(5):801–810.

32. Thames AD, Mahmood Z, Burggren AC, Karimian A, Kuhn TP. Combined effects of HIV and marijuana use on neurocognitive functioning and immune status. AIDS Care 2016;28(5):628–632.

33. Rizzo MD et al. HIV-infected cannabis users have lower circulating CD16+ monocytes and IFN-γ-inducible protein 10 levels compared with nonusing HIV patients. AIDS 2018;32(4):419–429.

34. Kallianpur KJ et al. Impact of Cannabis Use on Brain Structure and Function in Suppressed HIV Infection. J Behav Brain Sci 2020;10(8):344–370.

35. Manuzak JA et al. Heavy Cannabis Use Associated With Reduction in Activated and Inflammatory Immune Cell Frequencies in Antiretroviral Therapy–Treated Human Immunodeficiency Virus–Infected Individuals. Clin Infect Dis 2018;66(12):1872–1882.

36. Ellis RJ et al. Recent cannabis use in HIV is associated with reduced inflammatory markers in CSF and blood. Neurol Neuroimmunol Neuroinflamm 2020;7(5):e809.

37. Castro F de OF de et al. Distinct inflammatory profiles in HIV-infected individuals under antiretroviral therapy using cannabis, cocaine or cannabis plus cocaine. AIDS 2019;33(12):1831–1842.

38. Bredt BM et al. Short-term effects of cannabinoids on immune phenotype and function in HIV-1-infected patients. J Clin Pharmacol 2002;42(S1):82S–89S.

39. Krsak M et al. Self-Reported Cannabis Use and Markers of Inflammation in Men Who Have Sex With Men With and Without HIV. Cannabis Cannabinoid Res 2021;6(2):165–173.

40. Chao C et al. Recreational drug use and T lymphocyte subpopulations in HIV-uninfected and HIV-infected men. Drug and Alcohol Dependence 2008;94(1):165– 171.

41. Molina PE et al. Cannabinoid neuroimmune modulation of SIV disease. J Neuroimmune Pharmacol 2011;6(4):516–527.

42. Roth MD, Tashkin DP, Whittaker KM, Choi R, Baldwin GC. Tetrahydrocannabinol suppresses immune function and enhances HIV replication in the huPBL-SCID mouse. Life Sci. 2005;77(14):1711–1722.

43. Kieffer TL et al. G-->A hypermutation in protease and reverse transcriptase regions of human immunodeficiency virus type 1 residing in resting CD4+ T cells in vivo. J Virol 2005;79(3):1975–1980.

44. Bruner KM et al. Defective proviruses rapidly accumulate during acute HIV-1 infection. Nat Med 2016;22(9):1043–1049.

45. Bruner KM et al. A quantitative approach for measuring the reservoir of latent HIV-1 proviruses. Nature 2019;566(7742):120–125.

46. Bofill M et al. Increased numbers of primed activated CD8+CD38+CD45RO+ T cells predict the decline of CD4+ T cells in HIV-1-infected patients. AIDS 1996;10(8):827–834.

47. Benito JM et al. CD38 expression on CD8 T lymphocytes as a marker of residual virus replication in chronically HIV-infected patients receiving antiretroviral therapy. AIDS Res Hum Retroviruses 2004;20(2):227–233.

48. Rizzo MD et al. HIV-infected cannabis users have lower circulating CD16+ monocytes and IP-10 levels compared to non-using HIV patients. AIDS 2018;32(4):419–429.

49. Kumar V et al. Cannabinoid Attenuation of Intestinal Inflammation in Chronic SIV-Infected Rhesus Macaques Involves T Cell Modulation and Differential Expression of Micro-RNAs and Pro-inflammatory Genes. Front. Immunol. 2019;10. doi:10.3389/fimmu.2019.00914

50. Yin L et al. Anti-inflammatory effects of recreational marijuana in virally suppressed youth with HIV-1 are reversed by use of tobacco products in combination with marijuana. Retrovirology 2022;19(1):10.

51. Day CL et al. PD-1 expression on HIV-specific T cells is associated with T-cell exhaustion and disease progression. Nature 2006;443(7109):350–354.

52. Deeks SG. HIV Infection, Inflammation, Immunosenescence, and Aging. Annual Review of Medicine 2011;62(1):141–155.

53. Méndez-Lagares G et al. Specific patterns of CD4-associated immunosenescence in vertically HIV-infected subjects. Clinical Microbiology and Infection 2013;19(6):558–565.

54. Ibegbu CC et al. Expression of killer cell lectin-like receptor G1 on antigen-specific human CD8+ T lymphocytes during active, latent, and resolved infection and its relation with CD57. J Immunol 2005;174(10):6088–6094.

55. Tavenier J et al. Immunosenescence of the CD8(+) T cell compartment is associated with HIV-infection, but only weakly reflects age-related processes of adipose tissue, metabolism, and muscle in antiretroviral therapy-treated HIV-infected patients and controls. BMC Immunol 2015;16:72.

56. Streeck H et al. Epithelial adhesion molecules can inhibit HIV-1-specific CD8+ T-cell functions. Blood 2011;117(19):5112–5122.

57. Voehringer D, Koschella M, Pircher H. Lack of proliferative capacity of human effector and memory T cells expressing killer cell lectinlike receptor G1 (KLRG1). Blood 2002;100(10):3698–3702.

58. Wang S et al. An atlas of immune cell exhaustion in HIV-infected individuals revealed by single-cell transcriptomics. Emerg Microbes Infect 2020;9(1):2333–2347.

59. Eisenstein TK, Meissler JJ. Effects of Cannabinoids on T-cell Function and Resistance to Infection. J Neuroimmune Pharmacol 2015;10(2):204–216.

60. Xiong X et al. Cannabis suppresses antitumor immunity by inhibiting JAK/STAT signaling in T cells through CNR2. Signal Transduct Target Ther 2022;7(1):99.

61. Devi S et al. Immunosuppressive activity of non-psychoactive Cannabis sativa L. extract on the function of human T lymphocytes. Int Immunopharmacol 2022;103:108448.

62. Roth MD, Baldwin GC, Tashkin DP. Effects of delta-9-tetrahydrocannabinol on human immune function and host defense. Chem Phys Lipids 2002;121(1–2):229– 239.

63. Lin J, Zhong C, Hu D, Rudin C, Seltzer M. Generalized and Scalable Optimal Sparse Decision Trees. International Conference on Machine Learning. 2020

64. Maaten L van der, Hinton G. Visualizing Data using t-SNE. Journal of Machine Learning Research 2008;9(86):2579–2605.

65. McInnes L, Healy J, Saul N, Großberger L. UMAP: Uniform Manifold Approximation and Projection. Journal of Open Source Software 2018;3(29):861.

66. Wang Y, Huang H, Rudin C, Shaposhnik Y. Understanding How Dimension Reduction Tools Work: An Empirical Approach to Deciphering t-SNE, UMAP, TriMap, and PaCMAP for Data Visualization. Journal of Machine Learning Research. 2021; 73 (201): 1–73.

67. Chun T-W, Moir S, Fauci AS. HIV reservoirs as obstacles and opportunities for an HIV cure. Nat Immunol 2015;16(6):584–589.

68. Ipp H, Zemlin AE, Erasmus RT, Glashoff RH. Role of inflammation in HIV-1 disease progression and prognosis. Crit Rev Clin Lab Sci 2014;51(2):98–111.

69. Milloy M-J et al. High-intensity cannabis use associated with lower plasma human immunodeficiency virus-1 RNA viral load among recently infected people who use injection drugs. Drug Alcohol Rev 2015;34(2):135–140.

70. Chaillon A et al. Effect of Cannabis Use on Human Immunodeficiency Virus DNA During Suppressive Antiretroviral Therapy. Clinical Infectious Diseases 2020;70(1):140–143.

71. Abrams DI et al. Short-term effects of cannabinoids in patients with HIV-1 infection: a randomized, placebo-controlled clinical trial. Ann Intern Med 2003;139(4):258–266.

72. Kozela E et al. Cannabinoids decrease the th17 inflammatory autoimmune phenotype. J Neuroimmune Pharmacol 2013;8(5):1265–1276.

73. Howlett AC, Abood ME. CB1 & CB2 Receptor Pharmacology. Adv Pharmacol 2017;80:169–206.

74. Henriquez JE et al. Δ9-Tetrahydrocannabinold (THC) suppresses secretion of IFNα by plasmacytoid dendritic cells (pDC) from healthy and HIV-infected individuals. J Acquir Immune Defic Syndr 2017;75(5):588–596.

75. Henriquez JE, Rizzo MD, Crawford RB, Gulick P, Kaminski NE. Interferon-α– Mediated Activation of T Cells from Healthy and HIV-Infected Individuals Is Suppressed by Δ9-Tetrahydrocannabinol. J Pharmacol Exp Ther 2018;367(1):49– 58.

76. Alter G et al. HIV-1 adaptation to NK-cell-mediated immune pressure. Nature 2011;476(7358):96–100.

77. Harper J et al. IL-21 and IFNα therapy rescues terminally differentiated NK cells and limits SIV reservoir in ART-treated macaques. Nat Commun 2021;12(1):2866.

78. OngrÁdi J, Specter S, HorvÁth A, Friedman H. Combinedin vitro effect of marijuana and retrovirus on the activity of mouse natural killer cells. Pathol. Oncol. Res. 1998;4(3):191–199.

79. Healy ZR, Murdoch DM. OMIP-036: Co-inhibitory receptor (immune checkpoint) expression analysis in human T cell subsets. Cytometry A 2016;89(10):889–892.

80. Falcinelli SD et al. Longitudinal Dynamics of Intact HIV Proviral DNA and Outgrowth Virus Frequencies in a Cohort of Individuals Receiving Antiretroviral Therapy. J Infect Dis 2021;224(1):92–100.

